# TimeFlow 2: an unsupervised cell lineage detection method for flow cytometry data

**DOI:** 10.1101/2025.11.01.685988

**Authors:** Margarita Liarou, Thomas Matthes, Stéphane Marchand-Maillet

**Affiliations:** Department of Computer Science, University of Geneva, Switzerland; Hematology Service, Oncology Department, Hôpitaux Universitaires Genève; Clinical Pathology Service, Diagnostics Department, Hôpitaux Universitaires Genève; Centre Universitaire d’Informatique, University of Geneva, Switzerland

**Keywords:** trajectory inference, pseudotime analysis, flow cytometry, optimal transport, path clustering, cell differentiation, bone marrow data

## Abstract

Cell lineage detection refers to the inference of differentiation pathways from immature cells to distinct mature cell types. We developed TimeFlow 2, a new method for lineage inference in large flow cytometry datasets. It uses a single static snapshot of unordered cells and does not require prior knowledge of the number of pathways, cell type or temporal labels. TimeFlow 2 uses the cell orderings from TimeFlow and defines coarse cell states along pseudotime segments. By connecting these states, it constructs paths at cell state level. To approximate the trajectory structure, it further groups the paths based on an Optimal Transport-based cost function. We used TimeFlow 2 on three healthy bone marrow samples and accurately assigned monocytes, neutrophils, erythrocytes and B-cells of different maturation stages to four distinct pathways. Marker dynamics across the inferred pathways showed highly correlated patterns for the corresponding lineages in all three patients. We compared the performance of TimeFlow 2 and three other established methods using standard classification and correlation metrics. TimeFlow 2 outperformed the others on flow cytometry datasets and remained competitive on the challenging mass cytometry datasets. Overall, TimeFlow 2 detects biologically informative pathways, allowing bioinformaticians to model and compare marker dynamics across cell lineages in a data-driven way. Source code in Python and tutorials are available at https://github.com/MargaritaLiarou1/TimeFlow2.

## 1 Introduction

Trajectory inference (TI) methods infer cell lineage pathways formed during the dynamic process of cell differentiation. They compute a numeric value for each cell, known as pseudotime, based on similarities in cellular gene expression profiles, and use it to order the cells from the least to the most differentiated [1]. A characteristic application of TI methods is the process of hematopoiesis, or blood cell formation. This process takes place in the bone marrow and enables the production of functionally distinct blood cell types from hematopoietic stem and progenitor cells (HSCPs) [2]. TI methods have significantly contributed to the study of HSPCs fate decisions [3, 4, 5, 6], the study of hematopoietic gene regulatory networks [7], the comparison of healthy and dysregulated patterns, such as those observed in acute myeloid leukemia (AML) [8], and the analysis of gene expression alterations across different age groups [9]. Although most TI applications use RNA sequencing data (RNA-seq), their use has also gained interest in flow or mass cytometry (FC, MC). Examples of TI applications using FC or MC data include the reconstruction of healthy B-cell maturation trajectories and their comparison to those in patients with Wiskott-Aldrich syndrome [10], reconstruction of T-cell trajectories in HIV patients [11], and analysis of osteogenic lineage commitment during mesenchymal differentiation of skeletal stem cells [12].

The trajectory structure depends on the number of fully differentiated cell types involved in the process. Cells might progress towards a single mature cell type (linear trajectory), diverge along pathways that terminate in different cell types (tree-shaped and multi-branching trajectories) or evolve on a disconnected graph [13]. Different TI methods specialize in handling each case. We can also divide the methods by the form of the input data: either one static cell snapshot, assumed to capture all transitional states in the process, or multiple snapshots taken at different experimental time points (time-course datasets).

Many TI methods use a single static snapshot. They detect branches by constructing a piecewise linear Minimum Spanning Tree (MST) on cell clusters [14, 15, 16, 17, 18, 19, 20, 21, 22, 23] or a smoother MST with principal curves fitting [24, 25, 26] and principal manifold algorithms [27, 28, 29]. Alternatively, they represent cells with nodes on a k-Nearest Neighbour Graph (k-NNG) and simulate random walks to build an Absorbing Markov Chain, treating absorbing states as mature cell states [30, 31, 32, 33, 34, 35]. Other graph-based TI methods use heuristics such as correlation of random walks [36], measures such as graph node centralities [37], algorithms such as minimum cost flow [38], growing neural gas[39], community detection and statistical connectivity measures [40], hierarchical mixture modeling [41], and persistence homology diagrams for random walks clustering [42].

TI methods that require multiple snapshots with temporal labels model cell population dynamics assuming that cells differentiate in a least-cost way, and thus solve the discrete optimal transport (OT) problem to infer probabilistic couplings between cell populations in consecutive snapshots [43, 44, 45]. Methods that learn continuous cell dynamics from timecourse data use other OT frameworks, such as the dynamic OT formulation [46, 47, 48], the Jordan-Kinderlehrer-Otto flow scheme [49], and Schrödinger Bridges [50, 51, 52, 53]. However, these methods do not explicitly associate cells with lineage pathways. StationaryOT[54] and MultistageOT [55] propose OT couplings for a single static snapshot, but both require manually selected mature cells to infer the trajectory.

In this study, we focus on cell lineage detection in multi-branching trajectories using a single static snapshot obtained by flow cytometry. TI methods specifically designed for FC/MC data include Wanderlust [56] for linear trajectories, Wishbone [57] for single-branched trajectories, a method that adapts Wanderlust to tree-shaped trajectories, but requires manual specification of fully differentiated cells [58], CytoTree [59], which requires downsampling for pseudotime computation in large datasets, t-Space [60], and tviblindi[42]. Other related methods are TrackSOM [61] and ChronoClust [62], which track cell populations in timecourse datasets. To overcome challenges such as prior knowledge of terminal cell types or number of branches, downsampling and dependence on time-course data, we extend our previous method for pseudotime computation, TimeFlow [63], with an unsupervised cell lineage detection method for large cytometry datasets. Our method associates each cell with a pathway, allowing for certain cells to be shared across different pathways, reflecting their common ancestry. We demonstrate the potential of TimeFlow 2 in dissecting hematopoietic cell lineages using seven previously published bone marrow (BM) datasets, and discuss its usefulness in modeling the marker dynamics. We also compare our method with other well-known TI methods that detect cell lineages automatically, and interpret the biological relevance of the results based on ground truth gating labels.

## 2 Methods and Materials

### 2.1 Problem definition

We begin by formalizing the goal of our lineage detection method and the setup. Given a marker expression matrix X = {*x_ij_*} ∈ *R^N^*^x^*^D^*, where each row corresponds to a cell *i*, with *i* ∈ [1*, .., N*] and each column to a flow cytometry marker such as cluster of differentiation (CD), surface or intracellular marker *j*, with *j* ∈ [1*, …, D*], our goal is to associate each cell with a pseudotime value in [0,1] and a lineage pathway. The expected output is a set of tuples, with each tuple corresponding to an inferred pathway. Each tuple includes the indices of the cells that make up a particular pathway, along with the pseudotime of these cells. Formally, we denote a tuple as T = {(*z_i_, t_i_*) | *i* = 1*, . . ., L*}, where *z_i_* the index of the *i*-th cell, *t_i_* its pseudotime, and *L* ∈ *Z*^+^ the total number of cells in the pathway. This is a straightforward output to fit the marker expression as a function of pseudotime per pathway. As previously stated, some cells might be assigned in more than one pathways (e.g., multi-potent cells that have not committed towards a specific lineage). The number of pathways is not fixed in advance, but rather determined automatically by the method.

### 2.2 Method overview

TimeFlow [63] addresses the cell ordering task. It assigns every cell a pseudotime value by computing on a k-NNG the density-driven shortest path from the cell to the root of the trajectory (i.e., a specified stem cell with high CD34 expression). In this way, it preserves the continuum of cell transitions within local neighborhoods. However, pseudotemporal ordering is not enough to reconstruct lineages. In TimeFlow 2 we approximate the global trajectory structure with an approach that preserves (a) the ordering information from pseudotime, and (b) the biological information from the cell features (here markers). We account for (a) by dividing the pseudotime axis in several segments and defining coarse cell states within each. This is essential in understanding how these ordered cell states diverge into different branches along pseudotime. To satisfy (b) we assume that cells committed towards the same lineage differentiate through similar paths. Therefore, we construct paths at cell state level, which we then aggregate into biologically relevant groups based on their similarity. Despite having access only to a single cell snapshot, criteria (a) and (b) allow us to track the evolution of cell state paths simultaneously in pseudotime and marker space. More concretely, TimeFlow 2 takes the following steps:

1. Pseudotime segmentation: segment the pseudotime axis into equal-width, partly overlapping segments.
2. Cell clustering per segment: cluster cells within each segment. Unlike several other TI methods that force cells into clusters using the whole snapshot, here we limit clustering per segment to obtain ordered clusters or cell states.
3. Path construction: connect cell clusters between consecutive segments based on maximum cell overlap to construct paths at cell state level.
4. Path grouping: group paths based on an OT-based similarity metric to construct a trajectory backbone.
5. Trajectory refinement: refine the trajectory backbone.

Figure 1 provides the schematic overview of TimeFlow 2.

**Figure 1:**
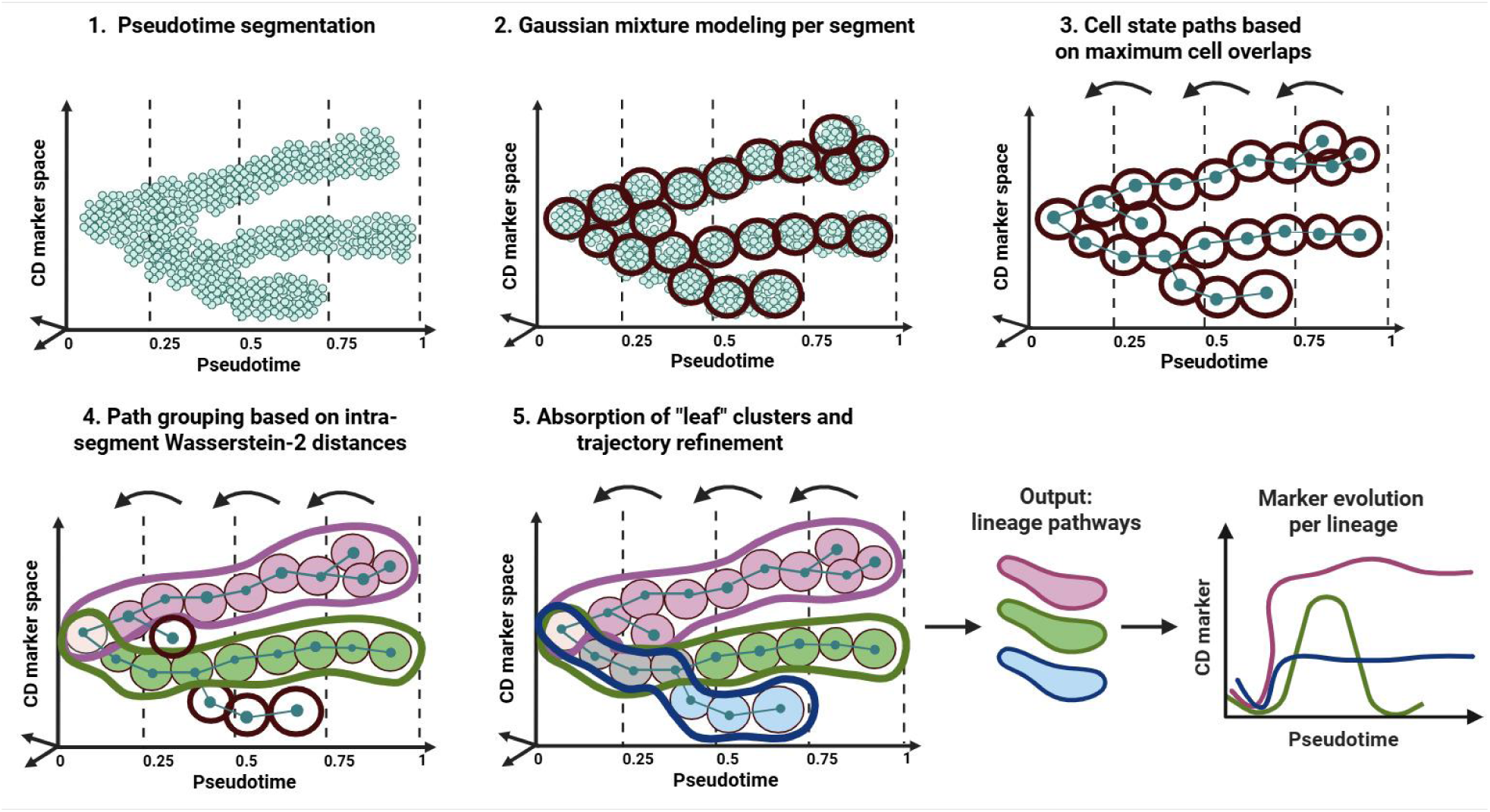
Schematic overview of TimeFlow 2. TimeFlow 2 takes as input a marker expression matrix and pseudotime values as estimated by TimeFlow. It follows the next steps. Step 1: Segments the pseudotime axis in equal width, partly overlapping segments (by default 35%). The overlap between segments is not included in this Figure. Step 2: Finds cell clusters within each segment using the Gaussian Mixture Model. Step 3: Connects clusters at consecutive segments based on the number of shared cells and constructs paths at cell state level. Step 4: Makes groups of paths based on their similarity. To compute the similarity of two equal length paths, it sums the intra-segment 2-Wasserstein distances between their Gaussian components at corresponding segments. Step 5: It refines the pathway groups by absorbing cell states that have not yet been integrated into any pathway. TimeFlow 2 assigns cells in different lineage pathways and enables downstream tasks, such as modeling marker expression as a function of pseudotime within each inferred lineage. The illustration was created using elements from BioRender.com.

### 2.3 Method details

#### 2.3.1 Pseudotime segmentation

We divide the pseudotime axis in equal-width segments, such that each consecutive pair of segments overlaps by a percentage of their width. We allow for overlapping cells between segments to retain the continuum of transitions, avoid strict boundaries, and later guide cell state connections. Hence, the lower bound of a segment precedes the upper bound of its previous segment. We denote each pseudotime segment by *s_i_*, where *i* ∈ 1*, …, S*, and *S* corresponds to the total number of pseudotime segments. Let {X*_si_* }_1_*_≤i≤S_* be the collection of the cells in *R^D^* for all pseudotime segments. The default value for pseudotime segments is *S* = 10, and 35% for segment width overlap.

#### 2.3.2 Cell clustering per segment

We model the cell populations within each segment *s_i_* with a Gaussian mixture model (GMM) [64], assuming that each Gaussian component describes a coarse cell state. In cytometry, GMM and its variants are particularly useful in automatically gating/clustering immune cell types [65, 66, 67, 68, 69, 70, 71, 72]. After estimating the GMM for each X*_si_*, we obtain

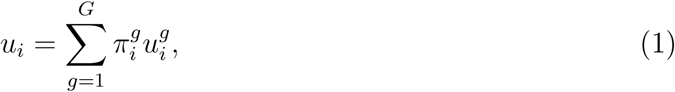

where *i* denotes the segment index, *G* denotes the number of total clusters in segment *i*, 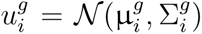 is a Gaussian distribution with a mean vector µ*^g^* and a covariance matrix Σ*_i_^g^* and π*_i_^g^* the weight of each Gaussian component for which 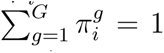 We note that each component has its own full covariance matrix and is characterized by its associated probability measure. We set *G* ≤ 25 ∀*s_i_*, and use the Bayesian Information Criterion (BIC) [73] to define the number of Gaussian components per segment.

#### 2.3.3 Path construction

In this step, we connect the clusters to define cell state transitions and construct paths at cell state level. Starting from the final segment, we connect each cluster in segment *s_i_* to a cluster in the previous segment *s_i−_*_1_ based on the maximum number of cells they share. We traverse pseudotime segments backwards to ensure that each cluster is connected to an earlier one, forming backtracked paths. Given the use of GMM for clustering, each path consists of Gaussian components across pseudotime segments and is denoted by

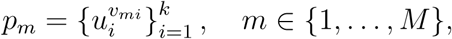

where *m* the path index, *M* the total number of paths, *u_i_* the full GMM model in segment *i*, *v_mi_* the specific Gaussian component at segment *i* that *p_m_* traverses and *k* the total number of segments *p_m_* traverses. To compute *M*, we count the total number of clusters across all segments and subtract from it the number of clusters in the first segment.

#### 2.3.4 Path grouping

To define a trajectory backbone, we aggregate paths that terminate at the final segment into biologically informative groups. We pose the question of how much effort is needed to transform one path into another. To quantify this we consider a cost function that sums the distances between the states of the paths at corresponding segments. We interpret the total cost as the degree of similarity between these paths. Small costs indicate similarity in the cell composition of the paths, and thus lineage agreement, while large costs imply differences in their cell composition and potentially lineage divergence. To measure the path similarity, we rely on OT. OT addresses the problem of mass transportation between two probability measures (or distributions) by coupling them via a transport plan based on a transportation cost function. OT defines a distance between probability measures when the transportation cost function is given by an *L^p^*-norm [74]. This is known as the p-Wasserstein distance, which describes the minimal cost of transporting the unit mass of one probability measure into the unit mass of the other measure. We provide more details on OT in Supplementary Section S1 and refer the reader to [75] for the theoretical foundations of OT, and to [74] for a computational-based approach. We represent the total path distance C between two paths as the sum of the 2-Wasserstein distances (*W*_2_) between their corresponding Gaussian components. For two paths *p_m_* and *p_m_′* of equal length *k* the distance is given as

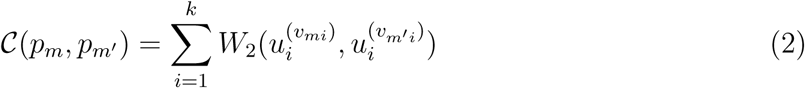

where 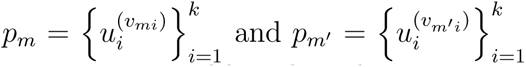 are two paths of same length *k*. Hence, the cost between two equal-length paths runs over all their segments *i* = 1*, . . ., k*, computing the *W*_2_ between their corresponding Gaussian components. Importantly, to compute these intra-segment *W*_2_ distances, we rely on a closed form solution that relieves us from computing full OT plans with *O*(*n*^3^) complexity. In the special case where both marginals *µ, ν* follow a (uni/multi-variate) Gaussian distribution (as is the case for *u_i_*^(^*^·^*^)^ here), *W_2_* can be explicitly written in terms of the mean vectors and the covariance matrices of the distributions, as noted in [76, 77]. For *µ* = N (µ_1_, Σ_1_) and *ν* = N (µ_2_, Σ_2_), the *L*^2^-Wasserstein distance is

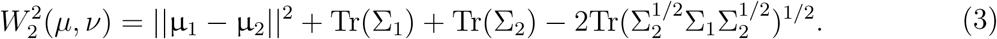

The pairwise path costs serve as inputs for unsupervised path grouping without access to the actual number of groups. Starting from no groups, we iterate over the path costs and merge two paths if their cost is below some threshold. If a pair of paths satisfies this condition but none of its paths belong to an existing group, a new group is created. Otherwise, if any path is already assigned to a group, the remaining path is merged into that group. It follows that paths whose costs from other paths exceed the threshold, will not be assigned to any existing group, but rather form their own path group. At the end of the above procedure, paths belonging to the same group receive a label (e.g, an integer). These labels are artificial and bear no biological information. They are used to indicate group memberships for the given threshold. Paths forming their own individual group also receive a label. Considering that choosing the cutoff threshold for the *W*_2_-based costs is not obvious, we repeat the same procedure for a range of candidate thresholds. The evaluation of this path grouping procedure is based on the model itself because no ground truth labels are available. We select as optimal threshold the one that maximizes the Silhouette score [78]. To compute SI, we use the implementation of [79] and give as inputs the artificial labels and the *W*_2_ costs. Higher thresholds for the *W*_2_ distances are more permissive for path merging compared to strict lower thresholds. To our understanding, the above pathway grouping strategy has not been used in the context of TI. An example where SI is not used for clustering validation, but rather for assessment of arbitrary groups of gene expression data is given in [80].

#### 2.3.5 Trajectory refinement

In the next step, we integrate in the trajectory backbone all the cell state paths that do not terminate at the final segment. These nuanced paths might be spurious or describe maturation towards new terminal states. Consequently, they are either absorbed by a previously discovered pathway or treated as new candidate pathways in the backbone. To identify such paths we search for clusters in any but the final segment with no forward connections. For simplicity, we refer to these terminal clusters as ”leaf clusters”. In addition, we list all clusters that belong to any of the main pathways and refer to them as ”explored clusters”. We distinguish the following cases for paths ending to a leaf cluster:

1. Paths corresponding to undiscovered pathways whose mature cells concentrate in segments earlier than the final one.
2. Paths corresponding to previously discovered pathways. This involves paths converging to a leaf cluster that is highly similar to an explored cluster either due to different differentiation speed or due to stochastic deviations from the main pathways. These deviations are highly likely considering that BIC tended to favor models with large number of clusters in step 2, especially in later segments. Moreover, the cell snapshot includes cells at different states of their own individual maturation, and thus variability and noise are expected in the trajectory.

To distinguish leaf clusters that mark potentially new terminal cell states from absorbable clusters, we use the intra-segment *W*_2_ distances, and examine which of the following cases is true for each leaf cluster:

1. None of the top-N (by default 4) closest clusters belongs to the list of explored clusters.
2. The top-1 closer cluster is an explored cluster but does not share highly correlated features with the leaf cluster.
3. The top-1 closer cluster is an explored cluster that shares highly correlated features with the leaf cluster.

On one hand, cases 1 and 2 motivate the inclusion of the leaf cluster’s path as a new main pathway. On the other hand, case 3 motivates the absorption of the leaf cluster’s path by one of the main pathways (after considering the update of the trajectory with new pathways). In case 1, leaf clusters may represent a previously unseen mature state. We require all top-4 closest clusters to be unexplored to reduce the risk of over-branching the trajectory, particularly in early segments. In case 2, we treat a leaf cluster that is not sufficiently correlated with its already explored top-1 closest cluster as a potentially new terminal state to prevent from under-branching the trajectory. In contrast, leaf clusters falling under case 3, as well as those that do not satisfy the condition of case 1, will be absorbed by an existing pathway. We treat the path of a leaf cluster as a time series, where the number of time steps equals the number of segments the path traverses. Each time step is characterized by a *D*-dimensional feature vector, with feature values computed based on the average marker expression of the cells falling in its clusters. We use the time series representation of each leaf cluster’s path to compute its Pearson correlation with the time series of its top-1 closest cluster and apply a strict threshold of 0.85 to distinguish between case 2 and 3. The threshold can be relaxed if the 10th percentile of all correlations across segments is above 0.85, indicating that even the weakest correlations maintain high similarity. Leaf clusters are absorbed by the existing pathways based on Fast Dynamic Time Warping (DTW) [81] which finds an optimal match between two multivariate time series independently of their length.

In the final step of our method we examine the hierarchical relationships between terminal cells from each pathway to mitigate the risk of spurious pathways. We retrieve the cells with largest pseudotime values per pathway (by default 5%) as representatives of their most mature cells, and label them by the pathway they belong to (pathway labels). We compute the marker summary statistics (mean, median, 25*^th^* and 75*^th^* quantiles) of those terminal cells, following other examples of feature engineering in cytometry [82], and proceed with hierarchical clustering. It follows that the threshold chosen to cut the tree hierarchy (dendrogram) drives the granularity of the clustering. A too low threshold might over-fragment the states, while a too high threshold might oversimplify the trajectory. Using the pathway labels (different from gating labels), we compute the Silhouette score at different thresholds and choose the one with maximum value (as described in 2.3.4). We note that our method allows users to vary this threshold based on their desired resolution. Optionally, users may consult relevant scatterplots of markers versus pseudotime for each pathway and experiment with different thresholds as explained in Section 3.5.

#### 2.3.6 TimeFlow 2 variant with FlowSOM

TimeFlow 2 is modular and allows replacement of its components for more flexibility. As an alternative to GMM clustering, we consider FlowSOM[83], which is a well-established non-parametric method for gating and visualizing cytometry data according to benchmarks[84, 85]. Similarly to Self-Organizing Maps (SOM)[86], FlowSOM trains a grid of SOM nodes and forms clusters by assigning each data point (cell) to its nearest node. Then, it updates the position of that SOM node and its neighbors to preserve the topological structure of the data. For easy integration with TimeFlow 2, which is written in Python 3.11.3, we choose the FlowSOM’s Python version[87]. We refer to this variant as TimeFlow 2 (FlowSOM) to distinguish it from TimeFlow 2 (GMM). Given that FlowSOM does not assume Gaussian distributions for the clusters, we cannot apply (3). We compute the intra-segment *W*_2_ distances of SOM nodes by solving the entropy-regularized OT problem[88] with the Sinkhorn’s algorithm for a fast and scalable approximation to OT plans (Supplementary Section S1). We set the default hyper-parameters of TimeFlow 2 (FlowSOM) to 10 segments, 35% segment overlap and 5x5 SOM grid (25 clusters) to match the default setting of TimeFlow 2 (GMM) and use a term of 1e-1 for entropic regularization and a batch size of 2048 for the OT plans. We discuss sensitivity to hyper-parameter settings for both variants in Section 3.3.

### 2.4 Datasets

We applied TimeFlow 2 on seven BM samples from patients with no hematological disease, as stated in the original publications. We re-used the P1/2/3-BM flow cytometry datasets from [63], which range from 499,544 to 597,613 cells and are characterized by 20 CD markers (Supplementary Section S1). Among cells at different maturation stages, we identified by gating 15%-20% mature neutrophils, 3%-6% mature monocytes, 0.3%-6% mature erythrocytes, and 0.7%-3% mature B-cells. All cell counts are given in Supplementary Table S1. In addition, we considered four other large, publicly available mass cytometry BM datasets. We used the Kimmey 6814 (347,167 cells) and Kimmey 6796 (647.730 cells) BM datasets, found in [89], which were annotated by the authors based on SPADE [90] clusters. Both datasets include 33 cell populations, among which the following cell types: neutrophils (23%), CD4 memory T-cells (8-9%), CD8 memory T-cells (8-9%), natural killers (NK, 7-10%), B-cells (4-6%), natural killers T cells (NKT, 1.1-3%), monocytes (1.1-2%), platelets (0.9-1.1%), conventional dendritic cells (cDCs, 0.9-1.6%), plasmacytoid dendritic cells (pDCs, 0.7%), basophils (Ba, 0.5%), polychromatophilic erythroblasts (0.2-0.6%), and plasma cells (0.1-0.4%). Supplementary Table S2 and Supplementary Table S3 summarize the cell counts for the Kimmey 6814/6796 datasets, respectively. Supplementary Section S2 lists the 32 antibody targets we retained for the analyses of these datasets. We also used the Levine 13 and Levine 32 datasets [91], which measure the cell expression with 13 and 32 CD markers in 167.044 and 265.627 cells, respectively. Levine 13 comprises of 25 cell populations, including mature B-cells (4%), monocytes (4%), mature CD4 T-cells (8%), erythroblasts (7%), mature CD8 T-cells (4%), NK cells (2%), megakaryocytes (2%), plasma cells (0.2%) pDCs (0.1%), platelets (0.003%, 5 cells), whereas Levine 32 contains 15 populations, out of which CD4 T-cells (9%), CD8 T-cells (7%), monocytes (7%), mature B-cells (6%), CD16 negative NK cells (1%), CD16 positive NK cells (0.8%), basophils (0.4%), pDCs (0.4%), and plasma cells (0.1%). We note that 50% of the Levine 13 cells and 60% of the Levine 32 cells remained uncharacterized and labeled as ”unassigned” in the original publication. Supplementary Section S2 provides more details on the panels used for these datasets, and Supplementary Table S4 and Supplementary Table S5 the cell fractions. Following standard preprocessing practices [92, 89], we transformed all MC datasets using the inverse hyperbolic sine (ArcSinh) with a cofactor of 5, and further scaled them to [0,1].

### 2.5 Evaluation metrics

To assess the biological relevance of lineages inferred by TimeFlow 2 and other TI methods, we used the cell type labels provided by the authors via manual or automated gating. Specifically, we defined reference lineages in line with the original publications and prior references to the literature of hematopoiesis [93]. For instance, in P1/2/3-BM datasets, we expect the following four lineages:

1. Monocytes: Immat Mono → Interm Mono → Mature Mono
2. Neutrophils: Immat Neu → Interm I Neu → Interm II Neu → Mature Neu
3. Erythrocytes: Immat Ery → Interm I Ery → Interm II Ery → Mature Ery
4. B-cells: Immat B-cells → Interm B-cells → Mature B-cells

Supplementary Section S3 provides the exhaustive list of expected lineages for the mass cytometry datasets we used (e.g., Monocytes: HSCs → Early Progenitors → Intermediate Progenitors → Late Progenitors → GMP → Myeloid → Monoblast → Pro-Monocyte → Monocytes).

These lineage references serve for the evaluation of the results with respect to 1) lineage purity, and 2) patterns of markers with known monotonically increasing expression in specific lineages. Lineage purity assesses the degree to which an inferred lineage includes cells associated with it. Patterns of lineage-specific markers indicate how faithfully a lineage reflects the expected increasing expression of certain CD markers. We compute the quantitative metrics as it follows:

1. Lineage purity: for each known mature cell type, we identified the inferred lineage pathway that contains the majority of its cells. We then defined as true positive cells (TP) those that are assigned to the identified lineage and truly belong to it based on the gating labels. False positive cells (FP) are those incorrectly included in the lineage, while false negative cells (FN) are those that should have been assigned to the lineage but were missed. To quantify the results we used the metrics of recall: TP/(TP+FN); precision TP/(TP+FP); and their harmonic mean F1-score: F1=2*TP/(2*TP+FP+FN). The F1-score ranges from 0 to 1, with higher values indicating both high precision and high recall. For instance, we label immature, intermediate, and mature B cells by *b*_1_*, b*_2_*, b*_3_, respectively, based on the gating labels, and similarly monocytes by *m*_1_*, m*_2_*, m*_3_, erythrocytes by *e*_1_*, e*_2_*, e*_3_*, e*_4_, and neutrophils by *n*_1_*, n*_2_*, n*_3_*, n*_4_. To evaluate the purity of the inferred B-cell pathway, we first identified the pathway that contained the majority of *b*_3_. In the same pathway, we then counted *b*_1_*, b*_2_*, b*_3_ that were correctly assigned to that pathway (TP). FP corresponds to the count of all other cell types that were included, and false negatives (FN), corresponding to *b*_1_*, b*_2_*, b*_3_ that were missing from the pathway.
2. Patterns of lineage-specific markers: as previously, we identified the inferred pathway that contains the majority of each known mature cell type. We selected well-studied canonical CD markers that increase monotonically within specific lineages and computed the Pearson, Spearman, and Kendall correlation between these markers and the pseudotime of cells assigned within that pathway. For instance, for pathways dominated by monocytes, we chose CD14; for B-cell pathways, we selected CD19 and CD20; and for neutrophils, CD16. Supplementary Section S3 gives the selected markers per dataset.

## 3 Results

### 3.1 Results on BM flow cytometry datasets

We first applied our method on the P1/2/3-BM datasets. Following pseudotime and lineage inference, we modeled the expression of selected markers as a function of pseudotime for each inferred pathway using a Generalized Additive Model (GAM) [94] with ten cubic splines. TimeFlow 2 inferred four pathways in the P1-BM dataset, which we further characterized by inspecting the evolution of lineage-specific markers in each of them, as shown in Figure 2A-P. CD14 expression was evident only for pathway 1 (Figure 2A) and CD16 expression rose only for pathway 2 (Figure 2F), implying that these pathways describe monocytic and neutrophilic differentiation, respectively. We note that CD14 smoothly increased along the pathway and the relative drop at pseudotime values close to 1 is attributed to outlier cells, as evidenced by the wider confidence interval bands near the end. CD36 expression increased in pathway 1 (Figure 2I) and showed a non-linear pattern in pathway 3 (Figure 2K), where it was initially upregulated for early cells, and then was smoothly downregulated for more mature cells. These CD36 patterns conform with monocytic and erythroid differentiation[95], respectively. Finally, only pathway 4 showed elevated levels for CD19 (Figure 2P), suggesting B-cell differentiation. Figure 2Q-T displays the cell distribution along pseudotime for each detected pathway, overlaid with the distribution of cells labeled with the corresponding ground truth trajectory. The histograms overlapped significantly and most cells spanned larger pseudotime values, which was in agreement with the larger counts of intermediate and mature cells in P1-BM. Pie charts in Figure 2U present the cell composition of each pathway across ten pseudotime segments of equal width. While the first segments of each pathway tended to contain a mixture of immature B-cells, neutrophils, monocytes and erythrocytes, later segments became more and more distinguishable and were dominated by a single mature population.

**Figure 2:**
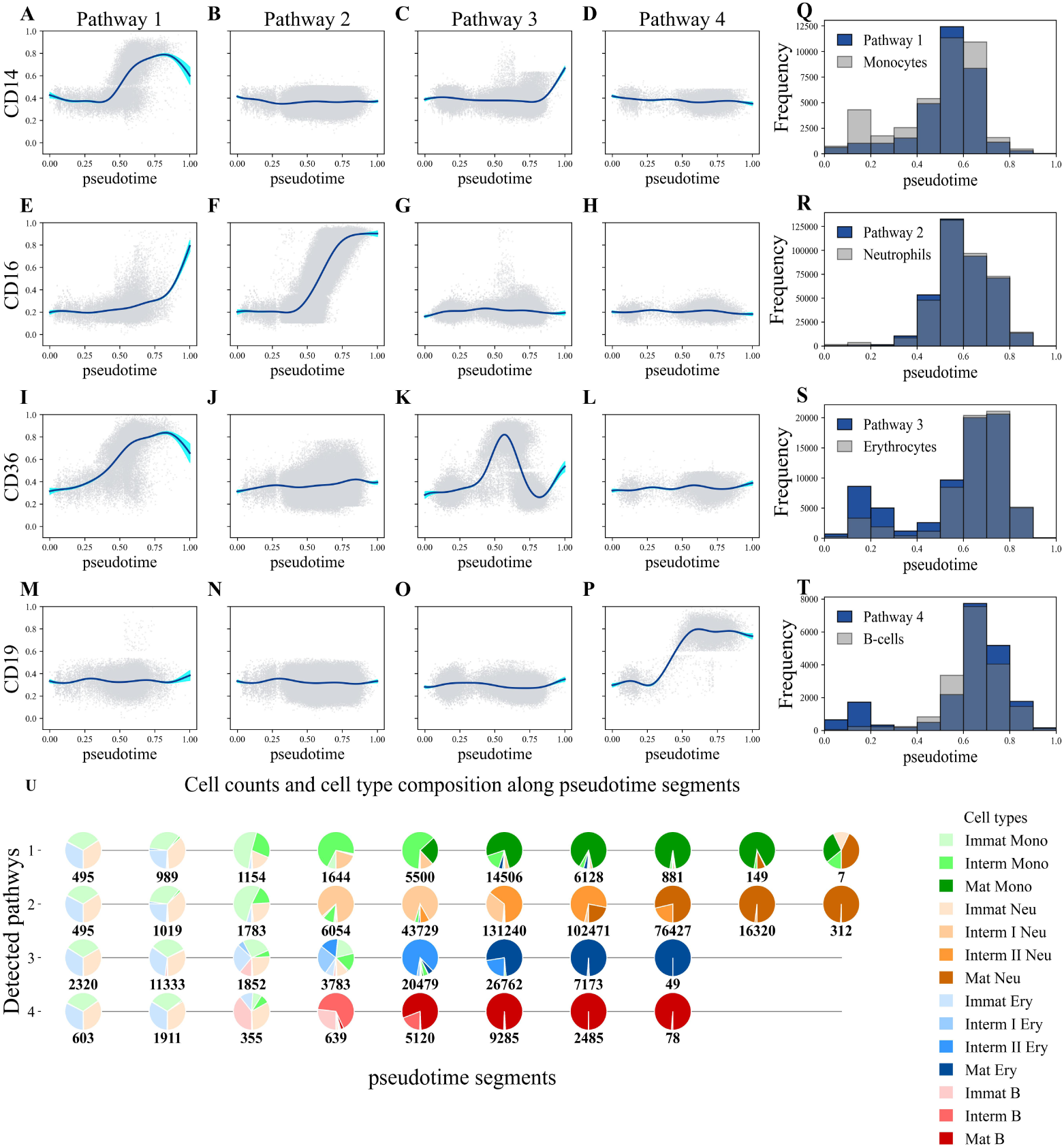
TimeFlow 2 results on the P1-BM dataset. Marker dynamics across the four different pathways/lineages inferred by TimeFlow 2. Each scatterplot dot represents a cell. Both pseudotime and marker expression are scaled in [0,1]. Solid blue curves show the Generalized Additive Model (GAM) fit used to model marker dynamics along pseudotime. Uncertainty around the estimated curve of each pathway is shown with shaded 95% confidence bands. (A-D) GAM fit of CD14 expression across the four pathways. (E-H) GAM fit of CD16 expression across the four pathways. (I-L) GAM fit of CD36 expression across the four pathways. (M-P) GAM fit of CD19 expression across the four pathways. (Q-T) Histograms with overlaid cell distributions along pseudotime. Dark blue histograms represent cell distri butions in the inferred pathways, while grey histograms reflect cell distribution of the known lineages based on the P1-BM ground truth gating labels. U) Pie charts showing the counts and cell type composition for each inferred pathway across ten pseudotime segments. Pie chart slices are colored based on the ground truth gating labels, with the total amount of cell counts noted for each pie chart.

All four populations share several cells prior to lineage commitment, which correspond to progenitor states not explicitly labeled in this dataset (e.g., HSCs, HSPCs). The heterogeneity of the early clusters may also be explained by the limited number of marker combinations used during manual gating, which resulted in substantial overlap in the distribution of several markers at the immature stage of these populations. As shown in Supplementary Figure S1, TimeFlow 2 (FlowSOM) also inferred four P1-BM pathways whose cell compositions matched accurately the monocytic (pathway 4), neutrophilic (pathway 1), erythroid (pathway 2) and B-cell lineages (pathway 3).

Our method inferred four pathways for the P2- and P3-BM datasets, too. Figure 3, presents 2D t-distributed stochastic neighbor embedding (t-SNE)[96] plots for the trajectories, allowing us to juxtapose the gating labels and the estimated pseudotime on the inferred pathways. The four distinct pathways clearly recovered the monocytic, neutrophilic, erythroid and B-cell differentiation, and shared some cells with lower pseudotime values. We emphasize that the t-SNE plots served only for visualization purposes. While several TI methods for RNA-seq apply dimensionality reduction or require 2D embeddings as input, we refrained from such steps prior to lineage inference to avoid potential information loss.

**Figure 3:**
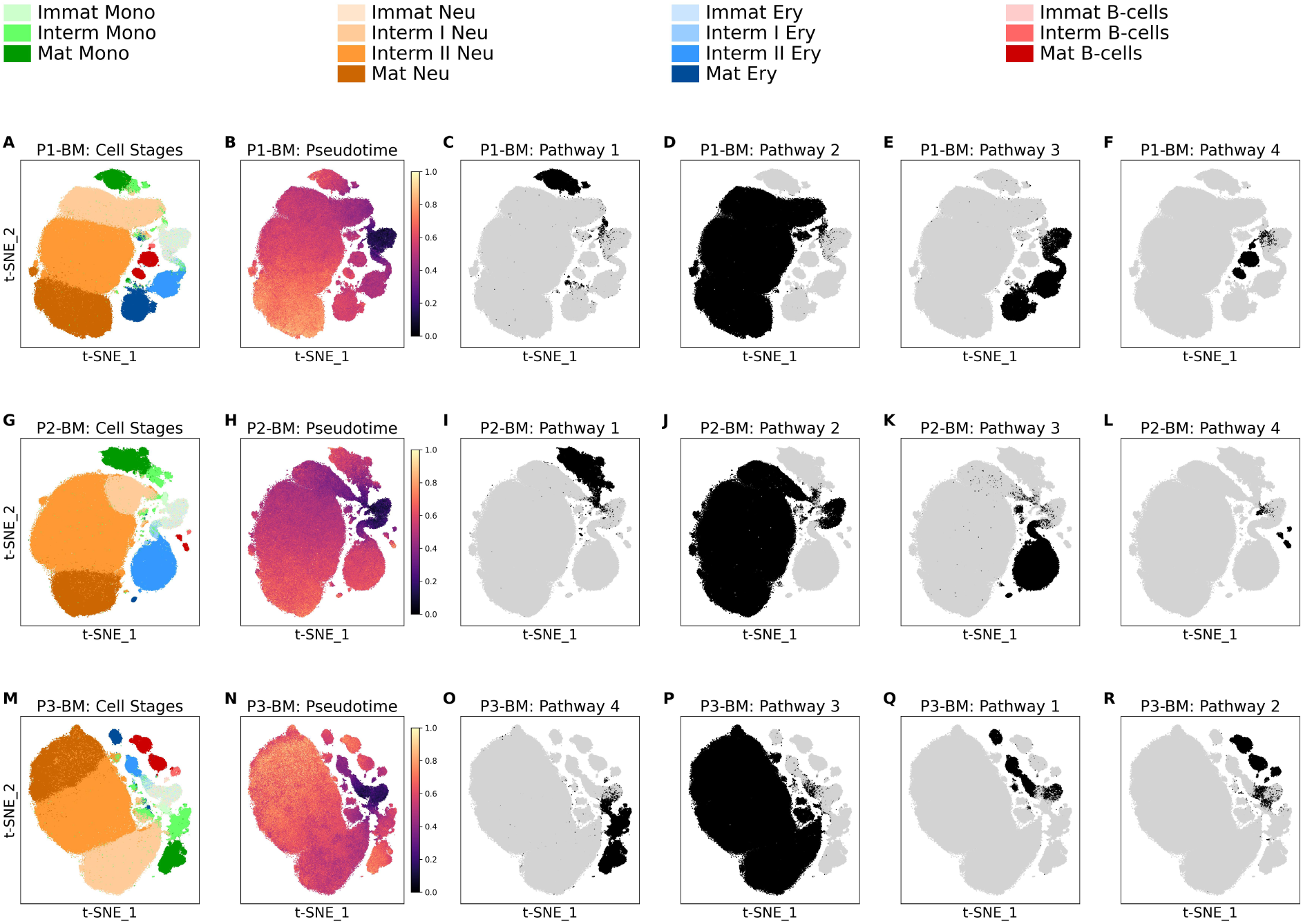
t-distributed Stochastic Neighbor Embedding (t-SNE) 2D projections of P1/2/3-BM datasets and pathways identified by TimeFlow 2. These t-SNE embeddings serve only visualization purposes and are not used during lineage inference by TimeFlow 2. Each row shows a t-SNE plot, where dots correspond to cells, colored 1) by their ground truth cell type and maturation stage, 2) by their pseudotime value, and 3) by their lineage pathway assignment as inferred by TimeFlow 2. (A-F) t-SNE plots for P1-BM. (G-L) t-SNE plots for P2-BM. (M-R) t-SNE plots for P3-BM. Inferred pathways are aligned vertically based on common lineages for easier comparisons across the patients.

We continued the analysis of P1-BM using Palantir [30] and VIA [31], two established TI methods for RNA-seq that are suitable for multi-branching trajectories, scale for large single cell datasets, identify automatically terminal cell states and assign each cell with some probability to reach any of these terminals (cell differentiation potential). We also considered CytoTree [59], which assigned cells to different branches of MST, but did not return pseudo-time values when applied on the full datasets. We did not include the well-known method of PAGA [40] in this comparison, because its graph output requires users to select manually both terminal nodes and nodes that comprise each lineage pathway. Moreover, we did not succeed in obtaining results for all patients using Slingshot [26] or Monocle 3[13] due to high memory demands. We supplied each method with the same root cells and followed their tutorials using the default hyper-parameters (Supplementary Section S4). Stacked barplots in Figure 4 present the cell type fractions and the total number of cells in each inferred pathway for each method. We observed that TimeFlow 2 consistently inferred four pathways and assigned the majority of cells from the same lineage into distinct pathways (Figure 4A,E,I). Palantir also detected four pathways for each patient, and in agreement with TimeFlow 2, it allocated immature cells from all populations across all pathways (Figure 4B,F,J). However, in the P1-BM dataset it distributed a considerable amount of intermediate II and mature erythrocytes in two distinct pathways (path 2 and path 3), and did not resolve the monocytic differentiation (Figure 4B). We observed that several cells such as mature monocytes were not assigned to any pathway, and thus tested a different strategy to map cells along the four detected branches based on their cell fate probabilities (Supplementary Section S4). CytoTree identified six lineages in P1-BM (Figure 4C), splitting most intermediate II neutrophils from their mature counterparts. Despite concentrating most B-cells into path 3, we noticed contamination with a significant proportion of mature erythrocytes and intermediate II neutrophils. VIA fragmented the trajectory into eight lineages ranging from 4,093 to 186,007 cells (Figure 4D). These included spurious pathways, such as path 2 (4,093 cells), which contained only intermediate I and mature neutrophils, and path 7 that attracted less than 5% of the total mature erythrocytes along with intermediate monocytes and neutrophils, but no other cells particular to the erythroid lineage. This indicates that further hyper-parameter tuning is needed (Supplementary Section S4). TimeFlow 2 and Palantir grouped cells in biologically meaningful pathways in both P2- and P3-BM, whereas CytoTree and VIA tended to disperse mature cells from the same lineage in different pathways (Figure 4G,H,K,L). Pathways obtained for P1- and P3-BM by TimeFlow 2 (FlowSOM) represented accurately the four anticipated lineages, while several of the P2-BM pathways were spurious, including mainly early cells (Supplementary Figure S2). Supplementary Figures S3-S5 present comparisons of marker dynamics across the inferred pathways of P1/2/3-BM for all TI methods. We noticed that the fragmented pathways of TimeFlow 2 (FlowSOM) for P2-BM did not distort the expected patterns because there were no unexpected mixtures of mature populations (Supplementary Figure S4E-H).

**Figure 4:**
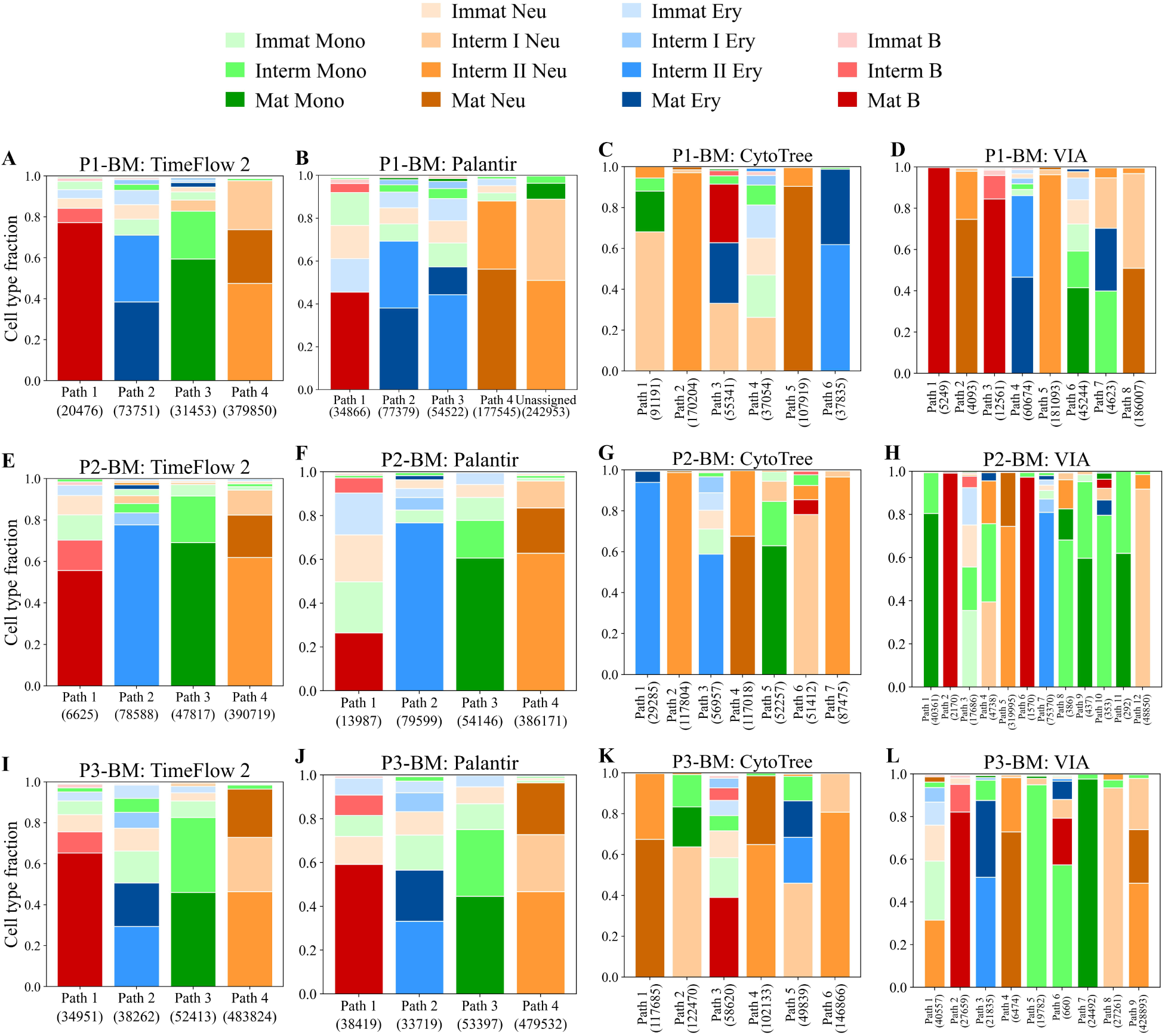
Side-by-side comparisons of cell type fractions across the different lineages identified by TimeFlow 2, Palantir, CytoTree, and VIA in the P1/2/3-BM datasets. Stacked bar plots show the cell distribution (%) in each inferred lineage path. The total number of cells assigned to each path is also given. Within each bar, cell types are presented by decreasing proportion. Cell types with larger percentages appear at the base of the bar, while those with smaller percentages appear at the top. (A-D) Bar plots for P1-BM dataset. (E-H) Bar plots for P2-BM dataset. (I-L) Bar plots for P3-BM dataset.

The above observations are also verified in Figure 5, which illustrates the evolution of CD14, CD16, CD36 and CD19 in the inferred pathways that contained the majority of mature monocytes, neutrophils, erythrocytes, and B-cells, for each patient and method, respectively. TimeFlow 2 variants resulted in coherent, smoothly increasing patterns for the expression of CD14 in monocytes, CD16 in neutrophils and CD19 in B-cells (Figure 5A,B,E,F,M,N) for all three patients. CD36 patterns were also highly correlated for P1- and P3-BM (Figure 5I,J). Despite not perfectly resolving the transitions of erythrocytes in P2-BM (Supplementary Figure S4C,G), both TimeFlow 2 variants captured the expected initial rise and later fall of CD36. Palantir confirmed the increase in CD16 expression for neutrophils and CD19 for B-cells, but missed the increase in CD14 for monocytes P1-BM (Figure 5C), as well as the nonlinear pattern of CD36 in P2-BM (Figure 5G, Supplementary Figure S4K). We attributed Palantir’s sharply increasing curves for CD14 in P2/3-BM monocytic pathways to the accumulation of numerous mature cells within too narrow pseudotime ranges (Supplementary Figures S4I,S5I) as shown by the barely visible confidence bands (95%) in Figure 5C. Supplementary Figures S3-S5 show similar disruptions in the continuum of transitions for several VIA scatterplots, where multiple cells are assigned the same pseudotime. The fragmented VIA trajectories affected mostly monocytes for which we noticed an unexpected decrease in CD14 expression (Figure 5D), and partially erythrocytes in P3-BM (Figure 5L), and B-cells in P2-BM (Supplementary Figure S4P).

**Figure 5:**
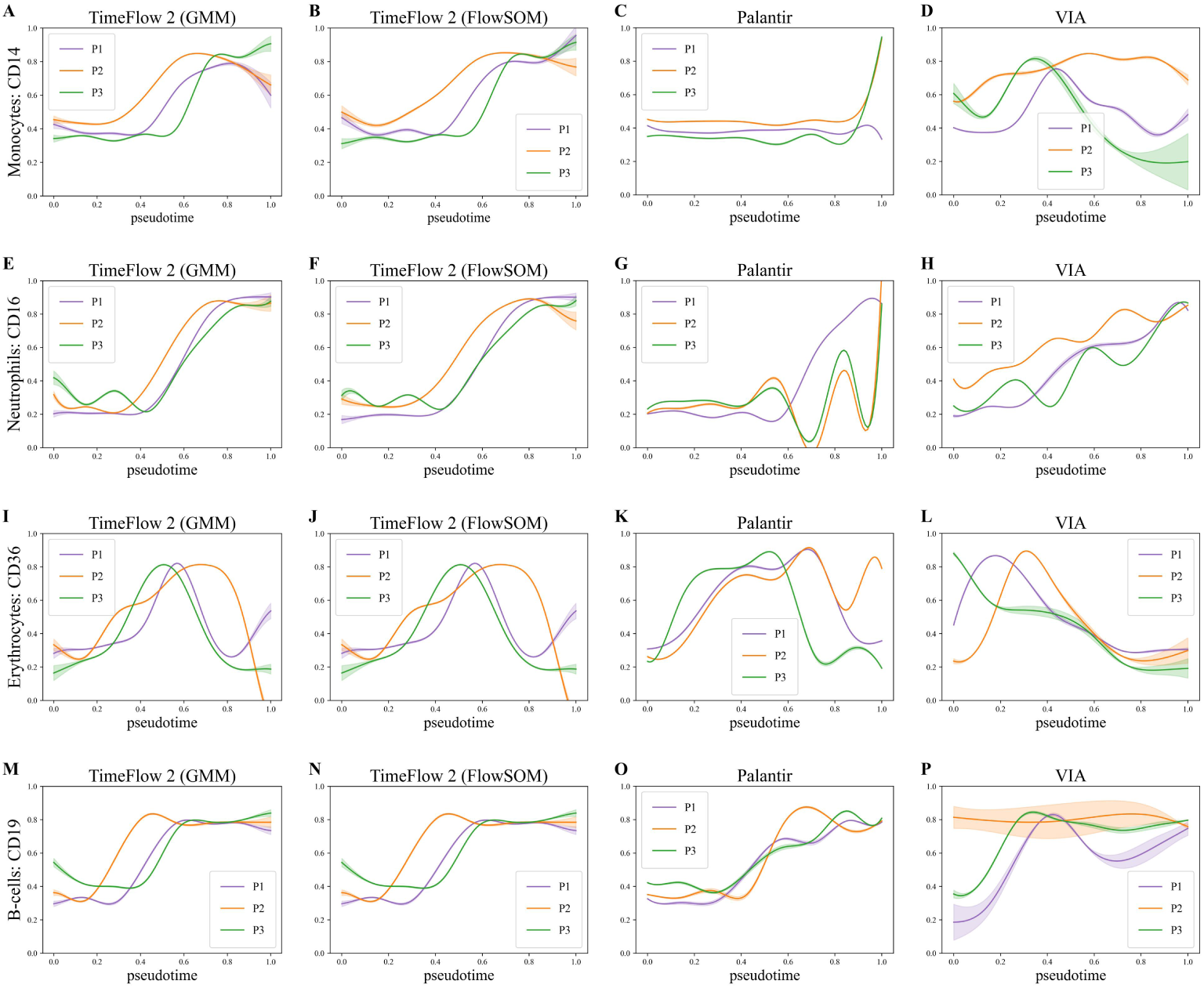
Side-by-side comparisons of marker dynamics across the different lineages identified by TimeFlow 2, Palantir, and VIA in the P1/2/3-BM datasets. None of the methods used gating labels during lineage inference. Labels were only used afterwards to retrieve lineages with most mature monocytes, neutrophils, erythrocytes, and B-cells. Both pseudotime and marker expression are scaled in [0,1]. Solid curves show the Generalized Additive Model (GAM) fit used to model marker dynamics along pseudotime and are colored by patient. Uncertainty around the estimated curve of each patient is shown with shaded 95% confidence bands. (A-C) GAM fit of CD14 expression across monocytes’ lineages. (D-F) GAM fit of CD16 expression across neutrophils’ lineages. (G-I) GAM fit of CD36 expression across erythrocytes’ lineages. (J-L) GAM fit of CD19 expression across B-cell lineages.

### 3.2 Results on BM mass cytometry datasets

We proceeded with the analyses of the public mass cytometry datasets. Before presenting the results, we remind the reader that pseudotime values might not always be biologically relevant [97]. Specifically, we note the case of T- and NK cells in BM datasets. Early T-cells, derived from the lymphoid lineage, migrate into the thymus to continue their development via double negative and double positive T-cells into CD4 and CD8 positive T-cells [98, 99, 100, 101]. This suggests that pseudotime for T-cell development should be estimated using thymus samples rather than bone marrow samples. Following the approach of other related studies [102, 103] using BM data, which did not include these populations in the differentiation hierarchy, we refrained from discussing expression patterns of T- and NK cells along pseudotime, and limited our assessment for these cell types to their grouping based on classification metrics. As noted in Supplementary Section S3, we did not consider early progenitors as part of the T-cell and NK reference lineages.

#### Kimmey 6814 dataset

TimeFlow 2 yielded eleven pathways for the Kimmey 6814 dataset and resolved accurately lineages of both major and rare populations. As shown in Figure 6A, it clearly separated mature and naive CD8 T-cells (pathway 1) from their CD4 counterparts (pathway 4). It assigned almost all NK cells in Pathway 11, and positioned the less abundant NKT cells (pathway 1) along with CD8 T-cells, without mixing them with myeloid-derived populations. We highlight pathway 7, where the rare population of basophils (0.05%) dominated, and pathway 8 which successfully absorbed the low abundant populations of pre-pDC and pDC (0.07%). Pathway 3 reflected differentiation of neutrophils as it mainly comprised of pro-myelocytes and mature neutrophils. Pathway 9 captured successfully erythroid populations such as MEP, pro-Erythroblasts, basophilic erythroblasts and polychromatic erythrocytes. B-cells were distributed in pathways 5 and 6, with the former containing about half as many cells as the latter, which showed a higher abundance of immature B-cells. We note that in Kimmey 6814 dataset, cells labeled as immature B-cells represent the terminal stage of the B-cell lineage (Supplementary Section S3). TimeFlow 2 did not separate monocytes (1.1%) from cDCs (1.6%), which were both assigned with myeloids. Supplementary Figure S6 shows the patterns of the lineage-specific markers in this dataset. TimeFlow 2 captured the expected increase in CD235a along the pathway where most erythrocytes were found, as well as the increase in CD123 and CD19 expression for the pathways where pDCs, and Immature B-cells prevailed (Supplementary Figure S6C,D,A). The presence of platelets in the monocytic/cDC lineage affected the quality of the GAM fit for CD14 (Supplementary Figure S6B,E). Palantir detected seven pathways (Figure 6B), offering good separation for T-cells, NK/NKT populations and neutrophils. However, it distributed polychromatophilic erythrocytes across all pathways, including those associated with lymphoid-derived populations. Likewise TimeFlow 2, Palantir also positioned platelets in the pathway where myeloid cells, cDCs and monocytes prevailed, displaying an inconsistent pattern for CD14 (Supplementary Figure S6L). Palantir assigned the rare population of basophils together with MEP, Pro-erythroblast, basophilic erythroblast and polychromatophilic erythrocytes. While early studies have reported GMPs as the origin of basophils [104, 105], later studies have suggested common progenitors shared between erythroid cells, megakaryocytes, eosinophils, mast cells and basophils [106, 107, 108]. Therefore, further experimental validation is needed. Results from CytoTree (Figure 6C) appear to be more convoluted, with T-cells and neutrophils suffering from fragmentation. Moreover, we observed that myeloid cells, cDCs and monocytes were assigned in pathway 3 along with B- and NK cells. VIA distinguished rare basophils from other populations (pathway 14), but overfragmented the pathways of NK cells, CD4 and CD8 T cells (Figure 6D). The decreasing expression in CD19 for the B-cell pathway 9 (Supplementary Figure S6P) indicated that the presence of T-cells across all pathways obscured the marker patterns.

**Figure 6:**
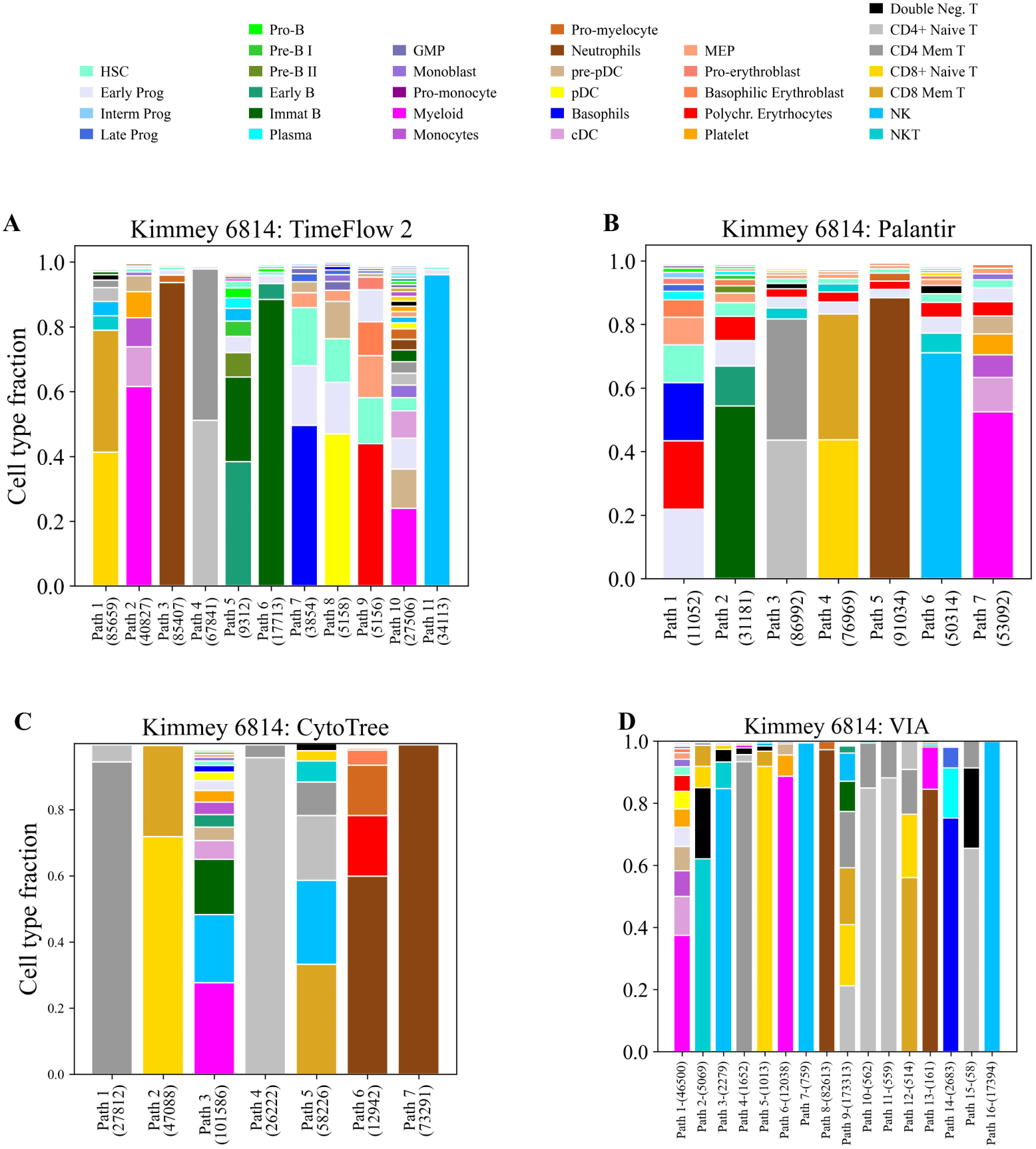
Side-by-side comparisons of cell type fractions across the different lineages identified by TimeFlow 2, Palantir, CytoTree, and VIA in the Kimmey 6814 dataset. Stacked bar plots show the cell distribution (%) in each inferred lineage path. The total number of cells assigned to each path is also given. Within each bar, cell types are presented by decreasing proportion. Cell types with larger percentages appear at the base of the bar, while those with smaller percentages appear at the top. (A) Bar plot for TimeFlow. (B) Bar plot for Palantir. (C) Bar plot for CytoTree. (D) Bar plot for VIA.

#### Kimmey 6796 dataset

TimeFlow 2 identified thirteen lineages in Kimmey 6796 (Supplementary Figure S7A), allocating in distinct pathways cells related to neutrophilic (pathway 9), erythroid (pathway 5), pDC (pathway 2), and B-cell differentiation (pathway 12). Moreover, it distinguished with high accuracy NK cells (pathway 8), CD4 T-cells (pathway 7) and plasma cells (pathway 11), but spread CD8 T-cells in pathways 1, 4, and 6. Monocytes (2.9%) were merged again with cDCs (0.9%) in pathway 3. Palantir detected five pathways and achieved good distinction among NK and CD4/CD8 T-cells (Supplementary Figure S7B). Surprisingly, it assigned a considerable proportion of myeloid cells and mature monocytes in every pathway, hindering insights from marker modeling for B-cell, monocytic, erythroid and pDC maturation (Supplementary Figure S8K-O). Furthermore, it associated pDCs with all five pathways and basophils with all but one pathway. CytoTree yielded less pure pathways in terms of CD4/CD8 T-cells (pathway 1) and merged several B-cells with myeloid-derived populations (myeloid cells, cDCs, pDCs, basophils) in pathway 7 (Supplementary Figure S7C). VIA (Supplementary Figure S7D) yielded 26 pathways, several of which were spurious considering that their sizes ranged from 69 to 106,074 cells. Lineage-specific expression patterns for Kimmey 6796 are illustrated in Supplementary Figure S8.

#### Levine 13 dataset

An observation common to each method for Levine 13 is that unassigned cells prevailed in almost all pathways (Supplementary Figure S9). TimeFlow 2 inferred ten pathways, with pathways 3, 5, 6 characterizing monocytic, erythroid/megakaryocytic and B-cell maturation, and pathways 8, 9, and 10 corresponding to NK, CD4 and CD8 T-cells (Supplementary Figure S9A). In the absence of erythrocyte-related markers in the Levine 13 panel, we highlight that TimeFlow 2, distinguished erythroblasts and megakaryocytes from other cell populations, but split them in two pathways (pathway 6 and 7). Moreover, it allocated several pre- and pro-B-cells in pathway 4, separately from the main B-cell pathway 5, making it less pure. Palantir found four pathways (Supplementary Figure S9B), separating accurately T-cell populations. However, mature monocytes were associated with each pathway, accounting for between 10.8% and 15.1% of each. CytoTree yielded seven pathways (Supplementary Figure S9C), and successfully accumulated in two independent pathways cells related with the erythroid (pathway 5) and monocytic differentiation (pathway 6). CD4 mature T-cells were split into pathways 4 and 7. Pathway 3 attracted mostly cells related to B-cell maturation, as well as NK cells. VIA inferred ten pathways (Supplementary Figure S9D). It unexpectedly merged populations such as MEP and erythroblasts with CD4 T-cells in pathway 2.

#### Levine 32 dataset

TimeFlow 2 detected seven pathways in Levine 32 (Supplementary Figure S10A). While it concentrated the majority of B-cells in pathway 3, and distinguished NK and NKT cells from T-cells in pathway 4, it did not resolve CD4 from CD8 T-cells (pathway 2). Furthermore, it produced two spurious branches for monocytes. Pathway 7 attracted underrepresented myeloid-derived cells (≤ 0.5%) such as basophils and pDCs, and pathway 6, which was the smallest, attracted mostly unassigned cells. Palantir successfully positioned monocytes, NK/NKT cells, CD4 and CD8 T-cells in four independent pathways (Supplementary Figure S10B). Likewise TimeFlow 2, Palantir placed pDCs together with basophils. B-cell maturation in pathway 1 was disrupted as several pre- and pro-B-cells were assigned in pathway 7. CytoTree inferred in total seven pathways (Supplementary Figure S10C), with pathway 4 consisting mainly of cells related to B-cell maturation. It fragmented monocytes into two distinct branches, and merged NK and NKT cells with CD8 mature T-cells in pathway 1. VIA suffered from over-fragmentation as it yielded 31 pathways that ranged from 10 to 46,183 cells (Supplementary Figure S10D).

#### Results with TimeFlow 2 (FlowSOM)

Supplementary Figure S11A shows the 14 pathways inferred by TimeFlow 2 (FlowSOM) for Kimmey 6814. The results showed clear separation between most CD8 and CD4 T-cells in pathways 1, 7, and prevalence of rare populations in distinct pathways, such as pDCs in pathway 2 and rare basophils in pathway 3. We also observed highly accurate recovery of erythroid differentiation in pathway 4, neutrophilic maturation in pathway 6, and dominance of NK cells in pathway 8. Unlike other methods, TimeFlow 2 (FlowSOM) successfully yielded a pathway specific to cDCs (pathway 14). However, the monocytic pathway 5 contained platelets, too. B-cells were spread across three pathways, with one of them including substantially more Immature B-cells, which represented the terminal stage for this lineage in this dataset. By using TimeFlow 2 (FlowSOM), we also obtained biologically faithful pathways in Kimmey 6796 (Supplementary Figure S11B), including the erythroid pathway 2, neutrophilic pathway 9, B-cell pathway 7, pDC pathway 12, as well as NK pathway 6, CD8 T-cells pathway 5, CD4 T-cells pathway 1, and plasma cells pathway 3. In Levine 13 (Supplementary Figure S11C) TimeFlow 2 (FlowSOM) performed strongly for several lineages. It assigned NK cells in pathway 5, CD8 and CD4 T-cells in pathways 7 and 6, respectively, and rare pDCs in pathway 9, but suffered from spreading in monocytes in pathways 1 and 14, and cells from the erythroid lineage in pathways 9 and 12. Results for Levine 32 were in high agreement with TimeFlow 2 (GMM) as shown in Supplementary Figure S11D.

Figure 7A-E summarizes the above findings by illustrating boxplots with the distribution of F1-scores for each method, first collectively for the P1/2/3-BM FC datasets, and then separately for the four MC datasets. Results are averaged across all expected lineages within each dataset as listed in Supplementary Section S3. Supplementary Figure S12 present the corresponding results for recall and precision. TimeFlow 2 variants had the highest F1 medians and lowest variability in P1/2/3-BM. TimeFlow 2 (GMM) had the second best median in Kimmey 6814, displaying lower interquartile range compared to other methods, with two outliers. In Kimmey 6796, its F1 median was the highest but the spread of the scores was larger compared to the previous dataset. We attributed the longer whiskers of all methods to their inefficiency in consistently dissecting less abundant cells such as cDCs. TimeFlow 2 (FlowSOM) outperformed other methods in Levine 13. Both TimeFlow 2 variants had lower medians compared to other methods in Levine 32. With regards to correlation metrics in Figure 7F-J, both TimeFlow 2 variants consistently exhibited lower variability compared to other methods, tended to have higher medians across all datasets and did not result in negative correlations, which would have implied significant distortions in the patterns of lineage-specific markers. As explained in [109], most methods originally designed for RNA-seq data, face severe scalability issues, often having quadratic or superquadratic complexity with respect to the number of cells and fail to analyze within an hour datasets of ten thousand cells. In terms of runtime, VIA was faster than the other methods completing full trajectory inference in less than one hour. TimeFlow 2 is one of the few TI methods that scales to large datasets within reasonable time. Steps 2.3.1, and 2.3.2-2.3.5 are executed instantaneously for TimeFlow 2 (GMM). Its burden is the clustering step. As shown in Supplementary Table S6, TimeFlow 2 (FlowSOM) consistently outperformed the GMM variant, completing lineage detection (clustering and OT plans) from 9 to 42 minutes when tested on datasets from 167,044 to 647,730 cells. Both variants were compared at their default hyper-parameters. However, the memory load of the GMM variant was lower due to the closed form solution for *W*_2_. Considering also the normalizing flow model trained for pseudotime computation[63], users should expect 20-90 minutes to analyze datasets of approximately 150,000-600,000 cells with TimeFlow 2 (FlowSOM) and 20 minutes to 2 hours for the same range of cells with TimeFlow 2 (GMM). In case of multiple healthy samples, users might consider sharing the normalizing flow weights across patients to speed up the analysis as discussed in [63]. We did not use GPUs for the normalizing flow models, which can significantly improve the runtime for full TI with TimeFlow 2.

**Figure 7:**
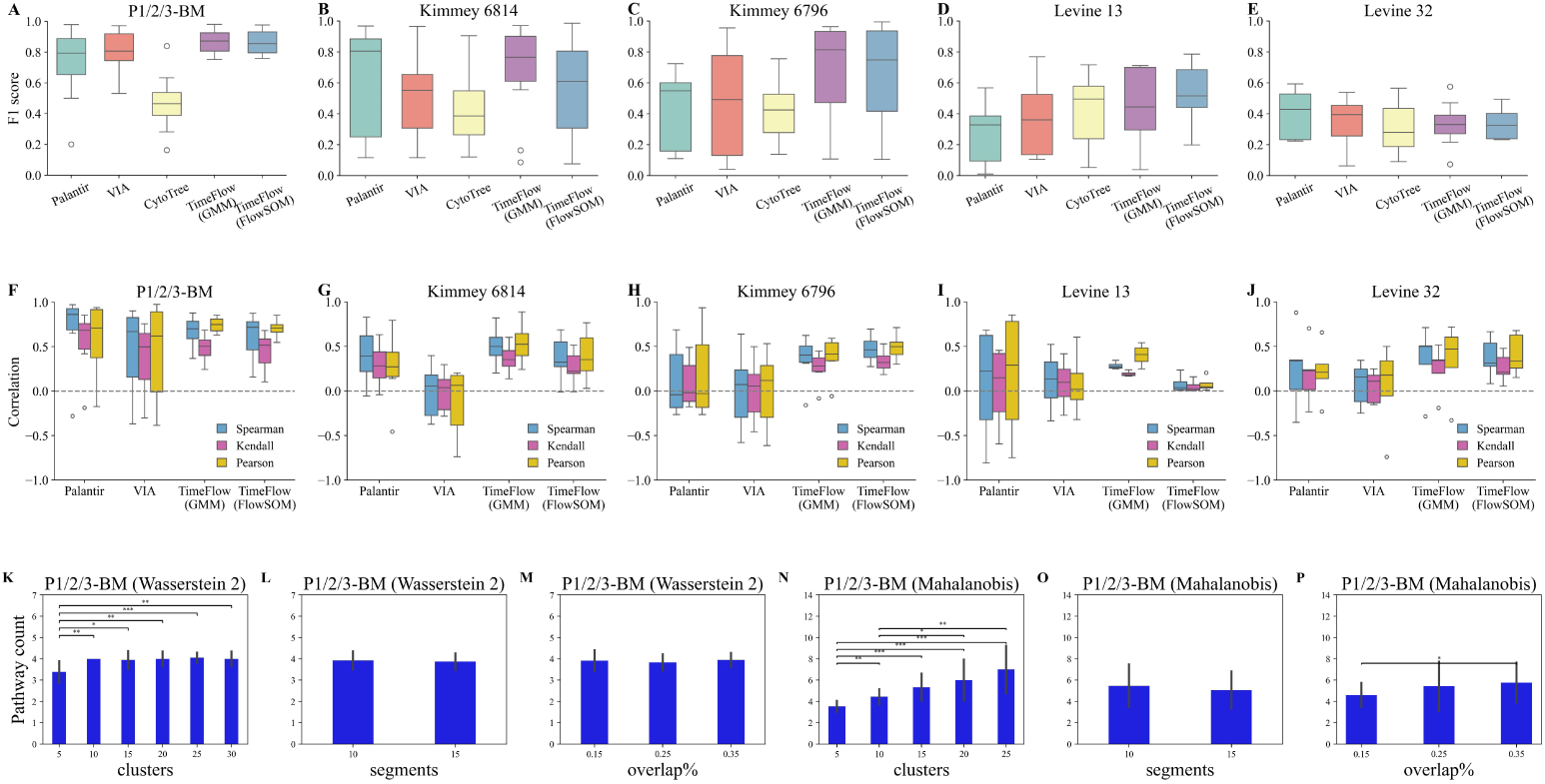
Methods comparisons across different datasets based on their F1-scores for all ground truth lineages identified in a dataset. Boxplots show the median, quartiles, minimum/maximum values, and outliers represented as individual points. (A-E) Results for the F1-score metric. (F-J) Results for the Spearman’s Rank, Kendall’s Tau and Pearson’s correlation coefficient between pseudotime in inferred lineages and lineage-specific CD markers. The horizontal dashed grey line at y=0 separates positive from negative correlation values. (K-P) Comparison between TimeFlow 2 using path costs based on the 2-Wasserstein metric versus a Mahalanobis-inspired metric. Each barplot shows the count of detected pathways over all P1/2/3-BM datasets for different hyper-parameter configurations. Error bars correspond to 95% confidence intervals. Benjamini & Hochberg adjusted p values correspond to paired one-sided t tests with a significance level of 0.05 (* signifies p value *<* 0.05, **: p value *<* 0.01, ***: p value *<* 0.001). Ground truth lineages in each dataset are described in Section 2.5 and Supplementary Section S3.

Overall, TimeFlow 2 yielded high scores for all classification metrics when applied on FC datasets, delineating accurately the myelomonocytic, lymphoid and erythroid lineages. It also resulted in coherent marker expression patterns across all three P1/2/3 patients. Regarding the more granular MC datasets, we found underrepresented mature cell types (0.05%-0.2%) to yield less stable results. For instance, both variants showed very good performance in Kimmey 6814 and 6796 datasets by detecting lineages where polychromatic erythroblasts, rare basophils and pDCs prevailed, but not in Levine 32. TimeFlow 2 remained competitive in terms of F1-score and as shown by the high correlation scores it did not violate the immunological hierarchies by mixing significant proportions of myeloid- and lymphoid-derived populations.

### 3.3 Hyper-parameter sensitivity and importance of *W*_2_

We applied both TimeFlow 2 variants on the P1/2/3-BM datasets using different configuration settings to test how their hyper-parameters affected the results. For TimeFlow 2 (GMM), we set the number of pseudotime segments to 10 or 15, the value of maximum number of GMM components per segment within the range [5,30] with incremental step of five, and the percentage of segment width overlap to 15/25/35%. For TimeFlow 2 (FlowSOM), we experimented with the same values of segments and overlaps, and set the SOM grid to 3x3, 4x4, 5x5, 6x6, which corresponded to 9, 16, 25, and 36 clusters per segment. These hyper-parameter combinations resulted in 2x6x3=36 and 2x4x3=24 experiments per patient for GMM and FlowSOM variants, respectively, which we first evaluated based on the number of detected pathways.

Figure 7K reveals sub-optimal performance for TimeFlow 2 (GMM) only when the number of maximum clusters per segment was five, implying that it is too low to resolve finely the cell states. We further evaluated the configurations in terms of F1-score for all cases. Supplementary Figure S13 presents a grid of F1-score boxplots, where each row corresponds to the maximum number of clusters, and each column to the segment width overlap. For each unique combination of those two parameters, we plotted the F1-scores against the number of segments. We observed that most boxplots contained 12 values, suggesting that TimeFlow 2 correctly recovered four pathways for all three patients. Clearly, the worst results were observed when the maximum number of GMM clusters per segment was equal to five (Supplementary Figure S13A-C). For all other configurations, median F1-scores remained close to 0.8 and had narrow interquartile ranges (Supplementary Figure S13D-R). Breaking down the results at population level (Supplementary Figure S14-S17), and focusing on configuration settings with more than five clusters, we observed high F1-scores and compact boxplots for monocytes (Supplementary Figure S14), and values close to one for neutrophils with practically no variability (Supplementary Figure S15). F1-scores remained close to 0.8 for erythrocytes (Supplementary Figure S16) with only four settings showing larger variability. Regarding B-cells, we noticed a drop in the F1-scores for several configurations (Supplementary Figure S17). We attributed this drop to the P2-BM B-cell lineage, which contained approximately only 5,000 cells. Although recall was close to 1 for all settings, precision was affected by the absorption of early cells from other populations, which as discussed in Section 3.1 was expected and did not distort the B-cell dynamics.

As illustrated in Supplementary Figure S18, TimeFlow 2 (FlowSOM) was robust across the configurations and yielded high F1-scores with low variability, except for three settings with 15 segments. We noticed inconsistencies in the number of inferred pathways, and we attributed this to spurious pathways we retrieved mainly for P3-BM. For all the previously mentioned configurations, we experimented with the Kimmey 6814/6796 and Levine 13/32 datasets using both TimeFlow 2 variants. We discussed in Supplementary Section S5 the robustness and variation in F1-scores between these two methods across all cell populations included in the MC datasets, and presented the boxplots in Supplementary Figures S19-S20. Overall, both TimeFlow 2 variants benefited from a larger number of clusters per segment when the datasets included more nuanced populations and a larger number of lineages was expected. We advise users to avoid reducing the number of segments as coarser segmentation might impact the discrimination of cell states in pseudotime. Results showed robustness and high median F1-scores for abundant populations such as neutrophils, B-cells, T-cells, and NK cells. Rare populations, occupying 0.05% to 2% of the total number of cells, yielded less stable results and posed a challenge in their detection for both variants. This might be attributed to lack of sufficiently discriminative markers and their small number rather than sensitivity to the hyper-parameter settings. As a good practice users may experiment with both TimeFlow 2 variants to construct an ensemble of lineage pathways, ensuring more robust and reliable cell lineage assignments.

In addition, we remark the importance of *W*_2_ in TimeFlow 2. We experimented with a different path cost function, inspired by the Mahalanobis distance (MD) [110] for all different hyper-parameter settings. We replaced equation (3) by 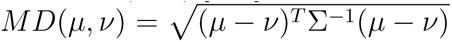, where *µ, ν* are two Gaussian components in our setup, and Σ a pooled covariance matrix given as the average of the full covariance matrices of the two components, 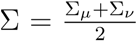. As shown in Figure 7 N-P, *W*_2_ consistently outperformed MD in terms of correct pathway number, suggesting that TimeFlow 2 yields more accurate and robust results with this distance. Unlike *W*_2_, MD tended to over-fragment the trajectory, and thus affected the F1 and correlation scores (not shown here). This observation supports the advantage of comparing cell distributions using their mean and covariance in a probabilistic way, derived via OT, instead of simpler centroid distances. More generally, other Wasserstein distances, such as *W*_1_ (also known as Mallow’s distance[111] or earth mover’s distance [112]), have proved useful in many computational methods for cytometry, including cell population and specimen comparisons [113, 114, 115, 116], evaluation of batch corrections [117], gating [118], dataset integration [119] and imaging cytometry [120].

### 3.4 Uncertainty of cell pseudotime and lineage assignments

The majority of TI methods provide a single point estimate of the pseudotime of each cell, ignoring its statistical uncertainty associated with high variability in gene/marker expression or technical sources [121]. However, single pseudotime estimates might bias lineage detection. We estimated the stability of pseudotime estimates with [63] by creating random subsets of each of the original P1/2/3-BM datasets using stratified sampling and maintaining 40% of the total number of cells (200,000-239,000 cells). For P2-BM, we created three ”cohorts” of ten subsets because several TI methods, including ours, tended to produce less accurate pseudotime values for erythrocytes, as mentioned in Section 3.1. This was also shown in Supplementary Figure S9 in TimeFlow [63]). We did not analyze all cohorts together to maintain a large number of common cells across the subsets. After computing the pseudotime and inferring the lineages for all subsets, we evaluated the results of each patient based on i) Pearson correlation for pseudotime of same cells across the subsets and Spearman, Kendall correlations for their rankings, 2) variance of pseudotime estimates, 3) lineage assignment consistency at a single cell level, and 4) F1-score for lineage detection.

As shown by the heatmaps of Supplementary Figure S21, we obtained very high correlations for P1-BM and P3-BM and slightly lower for the P2-BM cohorts. The pseudotime variance distribution of the common cells was heavily right-skewed for all patients, suggesting that most cells had very low variation. We also assessed the consistency of the lineage assignments for those common cells. Specifically, we examined whether the cell type composition of each cell’s assigned pathway(s) was consistent across the different experiments. We stored all the pathways in which each cell appeared and calculated the dominant cell type within each of them. We converted these counts first to percentages, and then to probabilities. For instance, if a cell appeared in 10% of the experiments in a pathway where the majority was mature neutrophils (n4), and 90% of the experiments in a pathway where the dominating population was mature monocytes (m3), we got the corresponding probabilities of 0.1 and 0.9. We used these probabilities to compute the entropy of the lineage assignments based on Shannon entropy, *H* = − ∑*_i_ p_i_* · log(*p_i_*). We expect cells with consistent assignments across the experiments to result in very low entropy scores (ideally zero), while cells with inconsistent assignments to yield a more spread distribution of entropy scores, indicating inconsistencies in their lineage assignment. As shown in Supplementary Figure S22 the majority of cell assignments resulted in zero entropy, indicating high agreement. Supplementary Figure S23 showed that most experiments yielded F1-score distributions with narrow interquartile ranges and very high median values. We identified one outlier for cohorts 1 and 2, and two for cohort 3 of P2-BM. By looking more thoroughly at the experiments which yielded these outliers, we found mature erythrocytes to be part of the neutrophilic lineage, and consequently leading to lower F1-scores. This aligned with our previous comment on the P2-BM erythrocytes, for which some mature cells received earlier than expected pseudotime. However, we note that when TimeFlow 2 used the full P2-BM, the misordered erythrocytes did not prevent us from recovering a pathway reflecting erythroid differentiation. This showed that TimeFlow 2, being able to process large cytometry datasets and operate independently from downsampling, protected us from these misordered cells.

In Supplementary Section S6, we discussed the impact of distorted pseudotemporal orderings on the capacity of our method to assign misordered cells in the expected lineages. Using the originally estimated pseudotime values of P1/2/3-BM, which led to biologically faithful lineages as shown in the Section 3.1, we created six new datasets per patient, each corresponding to a different amount of distortion on the original cell orderings. We used our method to find new lineages per dataset and evaluated the results with F1-score, Precision and Recall. Overall, TimeFlow 2 consistently yielded F1 medians above 0.8 and highly robust precision scores (Supplementary Figure S24). However, we observed a drop in the recall. From a biological perspective (and after verifying with the number of inferred pathways per case), we observed that as the amount of misordered cells increased, the method tended to overfragment the trajectories, producing redundant pathways. These pathways did not mix cells from different mature cell types, and continued to maintain cells from the different stages of a specific lineage (good precision), but at lower counts due to overbranching (lower recall). As explained above and in Supplementary Section S6, Supplementary Figures S21-S24 showed that TimeFlow 2 is resilient in accurate lineage detection under the presence of cells with inaccurate pseudotime and benefited from using large datasets. Nevertheless, these were preliminary experiments, and more rigorous analysis is needed to understand the degree at which inaccurate pseudotime deteriorates lineage detection.

### 3.5 Practical user guidance

We recommend practices for analyzing lineages inferred by TimeFlow 2 and tuning its hyperparameters in case of over-/underbranching. As a starting point, users can compute the cell count and other descriptive statistics for the markers of each lineage along pseudotime segments. This provides an overview of the lineages at a coarse level of cell clusters. In addition, we propose modeling the expression patterns of the markers (e.g., by fitting GAM models) to compare differences at finer resolution, prioritizing when available, lineage-specific markers that serve as a proxy to validate the results. Both practices may also help users detect potential over- or underbranching of the trajectory. An indication of spurious, noisy lineages is when a pathway contains an extremely low cell count, especially when a large proportion of it is shared cells with other pathways. In practice, this means that our method did not absorb this pathway by a more robust one. To control for overbranching, users may revisit the trajectory refinement step in Section 2.3.5 which is responsible for the introduction of new pathways that do not terminate in the final segment. This step is executed instantenously and gives users the possibility to immediately retrieve new results without performing clustering from scratch. Users can decrease the initially strict Pearson correlation threshold from 0.85 to a slightly lower value that will encourage the absorption of more leaf clusters by the trajectory backbone. To detect cases of underbranching users can inspect the confidence intervals (CIs) of the GAM fitted curves, and particularly cases with wide CIs. Unlike narrow CIs which suggest consistent marker expression within a pathway, wider CIs indicate uncertainty in the fitted model and heterogeneous cells within the same pathway (e.g., platelets mixed with monocytes in Supplementary Figure S6B,E). In this case, users may opt to isolate a subset of cells for further analysis or refrain from applying the post-hoc refinement step (Section 2.3.5). Skipping this step guarantees at least as many lineages as in the final output. In cases where more nuanced lineages are expected, users can also increase the number of clusters within segments. In the future we may focus on tuning the lineage resolution through regularization of lineage-specific markers.

## 4 Discussion

We developed TimeFlow 2 to automatically detect cell lineages in a single static snapshot. Our method builds on TimeFlow [63] for pseudotime computation. It operates independently from cell labels and prior knowledge of number of branches, enabling comparisons of marker dynamics across the trajectory in a data-driven way. TimeFlow 2 is modular and allows the replacement of GMM clustering and closed form solution for *W*_2_ by FlowSOM and entropic regularization of OT, respectively. By using using TimeFlow 2 on three large BM datasets obtained by FC, we confirmed cell differentiation into the myelomonocytic, erythroid and lymphoid lineages. Hematologists, biologists and bioinformaticians may use the pathway-specific scatterplots between CD markers and pseudotime to subsequently annotate the inferred lineages. They may also benefit from the flexibility of our method and guide the lineage dissection at their own desired resolution. We are interested in exploring further extensions of TimeFlow 2 for AML or myelodysplastic syndrome differentiation trajectories using flow cytometry data. Its use would bring an additional computational approach to current practices such as FlowSOM and predictive modeling in disease [122], and more generally clustering analysis for AML dynamics across disease stages [123].

Nevertheless, some challenges remain and require further investigation. Understanding more thoroughly the discriminative power of the CD markers and the relationship between their number and lineage resolution is important. This is motivated by the fact that Time-Flow 2 struggled separating CD4 and CD8 T-cells on Levine 32, but not on the Kimmey datasets. The detection of low abundant populations such as polychromatic erythroblasts, basophils and pDCs in some datasets was encouraging about the resolution TimeFlow 2 achieves. However, results for rare populations were less robust compared to major populations. It is also interesting to investigate other factors that may affect the detection of subtle pathways, such as discrepancies in manual or data-driven cell type labels across different studies, and the choice of clustering or dimensionality reduction methods. A further challenge in TI methods, as discussed in [30], is their common assumption on undirectional differentiation, where immature stem cells progressively differentiate into mature cell types. This is not necessarily true in diseased trajectories or cases of trans-/de-differentiation, where cells might return to previously acquired states. This requires caution when applying TI methods to complex dysregulated trajectories. Another challenge in TI is the lack of alignment between pseudotime and physical time [36]. To strengthen results obtained by TimeFlow 2 on a single cell snapshot, an interesting future direction involves comparisons to lineage inference and expression patterns from in-vitro experiments with access to multiple snapshots. This would allow us to inspect the timing of key events such as up-/downregulation of markers across lineages and make connections between pseudotime and real experimental time.

We also note the OT-based methods of LineageOT [124] and moslin [125], which take a different approach in TI by using RNA-seq time-course datasets that are not only equipped with experimental time points, but also with lineage information per point (e.g., via lineage tree and/or cell growth). In addition, GeneTrajectory [126] constructs gene instead of cell trajectories to account for other biological processes concurrent to cell differentiation. In the same spirit of validation using multiple datasets or single cell techniques, we are interested in consolidating results from TI analyses on flow cytometry data with those obtained using other technologies such as lineage clonal tracing [127, 128]. This would ensure further biological consistency of the inferred trajectories and more thorough understanding of (epi-)genetic programs that regulate cell differentiation.

## Code Availability

TimeFlow 2 is implemented with Python 3.11.3. The code is available on GitHub at https://github.com/MargaritaLiarou1/TimeFlow2. Interested users will also find tutorials for lineage inference and visualization with TimeFlow 2. For GMM, hierarchical clustering and SI computation we used the scikit-learn library (1.2.2)[79]. To compute clusters with FlowSOM we used its Python version (0.0.3)[87]. To approximate OT plans with Sinkhorn distances we used the POT Python library (0.9.1)[129]. We used pyGAM (0.8.0) [130] for fitting GAM curves.

## Data Availability

The public flow cytometry datasets P1/2/3-BM [63] are accessible in https://osf.io/ykue7/files. The public mass cytometry datasets Kimmey 6814 and Kimmey 6796 [89] are available in https://flowrepository.org/id/FR-FCM-ZYQV, while Levine 13 and Levine 32[91] can be downloaded from the R package HD-CytoData [131].

## Institutional Review Board Statement

The study was conducted in accordance with the Declaration of Helsinki and approved by the Institutional Ethics Committee of the University Hospital Geneva (Ethics Committee number 2020-00174 (date: 3 March 2020) and 2020-00176 (date: 28 April 2020)).

## Informed Consent Statement

Informed consent was obtained from all subjects involved in the study.

