## Supplementary Material for "TimeFlow 2: an unsupervised cell lineage detection method for flow cytometry data"

### 1 S1 Discrete OT and Wasserstein distance

Given the use of OT [1, 2] in steps 4 and 5 of our method, we briefly provide its essential background. We follow the Kantorovich formulation of OT, which addresses the problem of mass transportation between two probability measures (or distributions) by coupling them via a transport plan. Here, coupling refers to the joint distribution of these measures and the optimal plan is chosen based on the minimization of a predefined cost function. For two discrete probability measures, we consider two finite collections of points $\{x^{(i)}\}_{i=1}^{n}$, $\{y^{(j)}\}_{j=1}^{k}$, $x^{(i)}$, $y^{(j)} \in \mathbb{R}^D$, represented as empirical distributions: $\mu = \sum_{i=1}^{n} \mathrm{p}_i \delta_{x^{(i)}}$, $\nu = \sum_{j=1}^{k} \mathrm{q}_j \delta_{y^{(j)}}$, where $\delta_{x_i}$, $\delta_{y_j}$ are the Dirac delta functions centered at $x^{(i)}$ and $y^{(j)}$, respectively, with $x^{(i)}, y^{(j)} \in \mathbb{R}^D$ (these may also be considered as Dirac measures centered at $x^{(i)}$ and $y^{(j)}$). The corresponding probability masses are denoted by $p_i$ and $q_j$, such that the probability vectors p, q are non-negative and sum up to 1, i.e., $\sum_{i=1}^{n} p_i = \sum_{j=1}^{k} q_j = 1$. Given a positive cost function between pairs of points $c : \mathbb{R}^D \mathrm{x} \mathbb{R}^D \rightarrow \mathbb{R}^+$, OT seeks for a transport plan between $\mu$ and $\nu$ with minimal cost. Formally, it solves the following optimization problem:

$$OT(\mu, \nu) = min_{\Gamma \in \Pi(\mu,\nu)} < \Gamma, \mathrm{C} >= \sum_{ij} \Gamma_{ij} \mathrm{C}_{ij}, \quad (1)$$

where $\Gamma \in \mathbb{R}^{n\mathrm{x}k}$ is a coupling matrix whose entries $\Gamma_{ij}$ indicate the amount of mass flowing from $x^{(i)}$ to $y^{(j)}$, and $\mathrm{C}_{ij} = c(x^{(i)}, y^{(j)})$ is the cost matrix. Here <> represents the Frobenius dot product between matrices Γ and C. $\Gamma_{ij}$ is high if $x^{(i)}$ sends a lot of mass to $y^{(j)}$, and low otherwise. A transport plan $\Gamma^*$ is optimal if it achieves the minimum of equation (1). This optimization problem is subject to constraints $\Pi(\mu, \nu)$ that ensure Γ is measure-preserving, i.e., $\mu$ and $\nu$ are its marginals. The constraints are given as

$$\Pi(\mu, \nu) = \Gamma \in \mathbb{R}^{n\mathrm{x}k} | \Gamma 1 = \mathrm{p}, \Gamma^{\mathrm{T}} 1 = \mathrm{q}, \quad (2)$$

where 1 denotes a vector of ones. A nice feature of OT is that it defines a distance between probability measures when the transportation cost function is given by an $L^p$-norm [2]. This means that $\mathrm{OT}_c^p(\mu,\nu)^{1/p}$ is a distance metric between distributions (i.e., satisfies all metric axioms), with $c = L^p$ as cost function. It is called $p$-Wasserstein distance and describes the minimal cost of transporting the unit mass of one probability measure into the unit mass of the other measure. Specifically, the $p$-Wasserstein distance $W_p$ between $\mu$ and $\nu$ is defined as

$$W_p(\mu,\nu) = \mathrm{OT}_c^p(\mu,\nu)^{1/p}, \tag{3}$$

where $c$ the cost function and $p \geq 1$.

Problem 1 is a linear programming problem and its time complexity is $O(n^3)$. To avoid such complexity, [3] proposed a regularized OT objective as

$$OT(\mu,\nu) = min_{\Gamma\in\Pi(\mu,\nu)} < \Gamma, \mathrm{C} >= \sum_{ij} \Gamma_{ij}\mathrm{C}_{ij} + \epsilon \mathrm{H}(\Gamma), \tag{4}$$

where $H(\Gamma) = \sum \Gamma_{ij} log \Gamma_{ij}$ the entropy of the coupling matrix. 4 uses $H(\Gamma)$ as a regularizing function to obtain approximate solutions to 1. The objective in 4 is solved with the Sinkhorn-Knopp algortithm that provides an efficient and scalable approximation to OT. One downside of this regularization is that $OT(\mu,\mu) \neq 0$, which implies this quantity is no longer a valid distance. An alternative is the Debiased Sinkhorn divergence (not discussed here).

### 2 S2 CD marker panels

The panel of CD markers for the P1/2/3-BM flow cytometry datasets [4], includes the following 20 markers: CD200, CD45, CD45RA, CD64, CD3, CD15, CD133, CD117, CD56, HLA.DR, CD19, CD33, CD34, CD371, CD7, CD16, CD123, CD36, CD38. For both Kimmey 6814 and Kimmey 6796 analyses we selected 32 antibody targets that the original authors used to produce the SPADE clusters (see [5] Supplementary Table 4). We note a single difference in the dataset features: instead of including the anti-CD90-Biotin antibody, we used CD25. The full list includes: CD45, CD235ab, CD71, CD61, IgM, CD7, CD179b, CD11b, CD4, CD8, CD11c, CD34, CD179a, CD123, CD10, CD19, CD56, CD45RA, CD14, CD66, CD24, CD20, CD127, CD36, TdT, CD38, CD25, CD3, CD117, CD135, CD2, HLA.DR. The corresponding list for Levine 13 [6] is: CD45, CD45RA, CD19, CD11b, CD4, CD8, CD34, CD20, CD33, CD123, CD38, CD90, CD3, and for Levine 32 [6]: CD45RA, CD133, CD19, CD22, CD11b, CD4, CD8, CD34, Flt3, CD20, CXCR4, CD235ab, CD45, CD123, CD321, CD14, CD33, CD47, CD11c, CD7, CD15, CD16, CD44, CD38, CD13, CD3, CD61, CD117, CD49d, HLA-DR, CD64, CD41.

### 3 S3 Lineage references and markers used for evaluation

We describe each cell lineage and the cell populations used for calculation of F1-scores, precitions and recall (Section 2.5). We used the SPADE tree illustrated in Supplementary Figure 6 in [5] to define the transitions within each lineage.

**Lineages in Kimmey 6814 and Kimmey 6796**

1. B-cells: HSC → Early Progenitors → Intermediate Progenitors → Late Progenitors → Pro B-cells →Pre B I cells →Pre B II cells → Early Immature B cells → Immature B-cells
2. Monocytes: HSC → Early Progenitors → Intermediate Progenitors → Late Progenitors → GMP → Myeloid → Monoblast → Pro-Monocyte → Monocytes
3. pDCs: HSC → Early Progenitors → Intermediate Progenitors → Late Progenitors → GMP → pre-pDC → pDCs
4. Neutrophils: HSC → Early Progenitors → Intermediate Progenitors → Late Progenitors → GMP → Pro-Myelocyte → Neutrophils
5. Basophils: HSC → Early Progenitors → Intermediate Progenitors → Late Progenitors → GMP → Myeloid → Basophils
6. cDCs: HSC → Early Progenitors → Intermediate Progenitors → Late Progenitors → GMP → Myeloid → cDCs
7. Erythrocytes: HSC → Early Progenitors → Intermediate Progenitors → Late Progenitors → MEP → Pro-Erythroblast → Basophilic Erythroblast → Polychromatic Erythroblasts
8. Platelets: HSC → Early Progenitors → Intermediate Progenitors → Late Progenitors → MEP → Platelets
9. CD4 T-cells: Double Negative T-cells → CD4 Naive T-cells → CD4 Memory T-cells
10. CD8 T-cells: Double Negative T-cells → CD8 Naive T-cells → CD8 Memory T-cells
11. NK cells
12. NKT cells

**Lineages in Levine 13**

1. B-cells: HSC → MPP → Pre B-cells I → Pre B-cells II → Immature B-cells → Mature CD38 low B-cells → Mature CD38 mid B-cells
2. Monocytes: HSC → MPP → CMP → GMP → CD11b low Monocytes → CD11b mid Monocytes → CD11b high Monocytes

3. Erythrocytes: HSC → MPP → CMP → MEP → Erythroblasts

4. Megakaryocytes: HSC → MPP → CMP → MEP → Megakaryocytes, Platelets

5. pDCs: HSC → MPP → CMP → GMP → pDCs

6. CD4 T-cells: Naive CD4 T-cells → Mature CD4 T-cells

7. CD8 T-cells: Naive CD8 T-cells → Mature CD8 T-cells

8. NK cells

**Lineages in Levine 32**

1. B-cells: CD34+CD38- HSCs → CD34+CD38-CD123- HSPCs → CD34+CD38+CD123+ HSPCs → Pro B-cells → Pre B-cells → Mature B-cells

2. Monocytes: CD34+CD38- HSCs → CD34+CD38-CD123- HSPCs → CD34+CD38+CD123+ HSPCs → Monocytes

3. pDCs: CD34+CD38- HSCs → CD34+CD38-CD123- HSPCs → CD34+CD38+CD123+ HSPCs → pDCs

4. Basophils: CD34+CD38- HSCs → CD34+CD38-CD123- HSPCs → CD34+CD38+CD123+ HSPCs → Basophils

5. CD4 T-cells

6. CD8 T-cells

7. NK CD16 positive cells

8. NK CD16 negative cells

**Markers for evaluation**

To compute the correlation metrics, we selected well-known markers with increasing expression in specific lineages for each dataset. As explained in Section 3.2, we evaluated T-cells and natural killers only in terms of their lineage purity via the F1-score. We list below the markers used per dataset to evaluate the results.

**P1/2/3-BM**

- Monocytes: CD14
- Neutrophils: CD16
- B-cells: CD19

**Kimmey 6814 and Kimmey 6796**

- B-cells: CD19, CD20

- Monocytes: CD14
- Platelets: CD61
- Polychromatophilic erythroblasts: CD235ab

**Levine 13**

- B-cells: CD19, CD20
- Monocytes: CD11b (CD14 not available in the panel)
- pDCs: CD123

**Levine 32**

- B-cells: CD19, CD20
- Monocytes: CD14

### 4 S4 Methods comparisons

**Palantir:** We used Palantir [7] (palantir==1.3.0) following the tutorial https://palantir.readthedocs.io/en/latest/notebooks/Palantir_sample_notebook.html. To associate cells with the detected terminal states we used the function `select_branch_cells` with default values for q and eps. This function considers the differentiation potential/fate probabilities of the cells and their pseudotime, and returns an array of boolean masks (True/False) to indicate whether a cell belongs to the path of a terminal state. For the P1-BM dataset, we noticed that several cells were assigned a False value for all four detected terminal states. We did not manage to resolve the issue by tweaking the default values of q and eps, and labeled these cells as unassigned. Alternatively, we used the fate probability estimates (output from `palantir.core.run_palantir`), which returns the probability of each cell terminating in each of the four terminals, and picked the maximum numeric value for each cell. In this way, we ensured that every cell belonged to a lineage. Nevertheless, this did not improve the results for two reasons: 1) two of the identified terminal cells were mature erythrocytes, 2) by strictly mapping each cell to a unique lineage, we ended up with lineages including mainly mature cells (e.g., 100% mature erythrocytes) that did not represent the lineage evolution. We did not face this issue with P2/3- datasets. Our P1-BM analysis with Palantir is uploaded here: https://osf.io/u275b/?view_only=ed1b706952b949e88dd5de0da262f983. We did not find unassigned cells for P2/3-BM, Kimmey 6814/6796 and Levine 13/32 datasets.

**VIA:** We used VIA[8] (pyVIA==0.1.89) following the tutorial https://pyvia.readthedocs.io/en/latest/notebooks/ViaJupyter_scRNA_Hematopoiesis.html. Given the tendency of VIA to overfragment the trajectories, we experimented with some hyper-parameters related to the granularity of the results, such as `too_big_factor`, `neighboring_terminal_states_threshold`, and `resolution_parameter`. Although the default `resolution_parameter` is set to the maximum 1, (e.g., in time-course cytometry data https://pyvia.readthedocs.

io/en/latest/notebooks/mESC_timeseries.html), we found that among the previous hyper-parameters this one could decrease the number of branches. Nevertheless, we had to reduce the threshold to values below 0.5. However, we could not be conclusive, as the resolution threshold appeared to be sensitive within the same dataset, and same threshold values did not generalize across the three patients. For instance, by decreasing the resolution from 1 to 0.3 for P1-BM, we obtained four lineages that accurately separated the populations, but decreasing to 0.2 yielded two branches dominated by B-cells, a branch with mature neutrophils, and a branch with intermediate II neutrophils, mature erythrocytes and mature monocytes. By increasing the value to 0.4, we found seven branches that mixed several mature populations. We also set resolution to 0.3 in P2/3-BM, but found it insufficient to distinguish mature B-cells from neutrophils in P2-BM, and not suitable for merging mature neutrophils and mature erythrocytes with their ancestors in P3-BM. Our P1-BM analysis with VIA is uploaded here: https://osf.io/u275b/?view_only=ed1b706952b949e88dd5de0da262f983

**CytoTree:** We used CytoTree [9] following the tutorial https://ytdai.github.io/CytoTree/analysis.html#pseudotime. To extract the cell branch assignments we used the function `buildTree` with raw dimensions and the metadata property of the CytoTree object `meta.data$branch.id`.

### 5 S5 Hyper-parameter sensitivity for TimeFlow 2 variants

We used the hyper-parameter configurations presented in Section 3.3 to experiment with the Kimmey 6814/6796 and Levine 13/32 datasets for both TimeFlow variants and compare their F1-score distributions for all cell populations and discuss their robustness. Supplementary Figures S19-S20 present the relevant F1-score boxplots. We excluded from these boxplots all F1-scores for configurations with five clusters due to their insufficiency to resolve the four lineages in the P1/2/3-BM, as discussed in Section 3.3. In Kimmey 6814 (Supplementary Figure S19A-B), both TimeFlow 2 variants excelled in the detection of neutrophils, CD4, CD8 T-cells, and B-cells with F1-scores close or above 0.8 and narrow interquartile ranges. These populations along with the NK cells were the most abundant of this dataset. In sharp contrast, less abundant populations such as platelets and NKT yielded consistently poor F1-scores. TimeFlow 2 (FlowSOM) variant clearly outperformed the GMM one with respect to NK cells and produced less variable scores for monocytes, indicating better robustness across the configurations. By looking at the actual values of F1-scores for NK cells, we found that these outliers were exceptionally produced when the SOM grid was set to 5x5 and the cell overlap to 25%. Moderate results for Erythroblasts were sufficient for the dominance of this population in distinct lineages, and their analyses tended to benefit from a larger number of clusters. Supplementary Figure S19C-D confirmed the previous findings for accurate and robust detection of the major lymphoid and myeloid lineages. Platelets and NKT cells suffered again from low scores. Several populations exhibited more robust F1-scores with the FlowSOM variant. By looking at the actual configurations, we attributed the longer lower whiskers of NK scores with TimeFlow 2 (GMM) to experiments with at most 10 GMM clusters per segment. In Levine 13 (Supplementary Figure S20A-B), we found the FlowSOM variant to be more robust, yielding consistently higher F1 medians for all cell types and more compact boxplots with shorter whiskers. FlowSOM clearly benefited TimeFlow 2 with respect to T-cell populations and erythroblasts, demonstrating better and more robust results. F1-scores for Levine 32 (Supplementary Figure S20C-D) tended to be lower than the ones in the previous datasets. Results were robust for B-cells and monocytes.

### 6 S6 Pseudotime distortion and impact on lineage detection

In this section we briefly discuss the impact of distorted pseudotemporal orderings. Given the sensitivity of pseudotime estimation to dataset-specific characteristics, and the absence of ground truth, it is reasonable to examine the impact of distorted cell orderings on the capacity of TimeFlow 2 to assign misordered cells across different lineages. Using the originally estimated pseudotime values of P1/2/3-BM, which led to biologically faithful lineages based on Results Section 3, we created six new datasets per patient, each corresponding to a different amount of distortion on the original cell orderings. Specifically, we distorted the rankings of the cells by randomly swapping the pseudotime values of 5%, 10%, 15%, 20%, 25%, and 30% of the total amount of cells. For these experiments we retained all cells (no downsampling) and swapped the rankings of 25,000 to 179,000 cells for all three patients based on random selection of cells. This means that we changed the rankings of 25,000 to 179,000 cells for all three patients based on random selection of cells. Then, we run TimeFlow 2 on the modified datasets for two different configurations (10 segments, maximum 25 GMM clusters, 35% overlap, and 10 segments, maximum 20 GMM clusters, 35% overlap), which yielded high scores for all the evaluation metrics when tested on the original pseudotime values. We recomputed the evaluation metrics, and provided in Supplementary Figure S24 the distribution of the new scores for all cell types of all three patients. Median F1-scores remained above 0.8 and pairwise comparisons with Benjamini-Hochberg corrections for multiple testing showed no statistically significant differences. Nevertheless, we observed more outliers for the heavily distorted 30% dataset, and lower medians and larger variability for datasets with more than 20% distortion. Overall, the method was robust with respect to precision scores. However, as the recall scores indicate, all distorted datasets resulted in statistically significant different results from the original one, leading to lower medians and more variable scores. From a biological perspective (and after verifying with number of detected branches per case), the above results mean that as the amount of misordered cells increased, the method tends to overfragment the trajectories and produced redundant pathways. These pathways did not mix mature cell types and continued to maintain cells from the same lineage, but at lower cell counts (hence good precision but worse recall). TimeFlow 2's robust precision scores show its ability to separate the cells into relevant lineages without heavily mixing cells from different mature cell types under distorted orderings. However, as expected inaccurate pseudotime impacts negatively downstream tasks such as marker expression modeling within each inferred lineage, even when the lineage pathway correctly captures the expected cell populations.

### Supplementary Tables

| Patient | Cell Population | Immature | Intermediate I | Intermediate II | Mature | Total Count |
|---|---|---|---|---|---|---|
| P1-BM | Mono | 7,703 | 12,629 | - | 18,924 | 39,256 |
| | Neu | 6,705 | 92,575 | 180,567 | 99,858 | 379,705 |
| | Ery | 6,012 | 1,801 | 24,204 | 30,377 | 62,394 |
| | B-cells | 883 | 1,444 | - | 15,862 | 18,189 |
| | BM | - | - | - | - | **499,544** |
| P2-BM | Mono | 9,368 | 16,280 | - | 33,081 | 58,729 |
| | Neu | 5,320 | 48,227 | 242,539 | 80,014 | 376,100 |
| | Ery | 1,720 | 4,603 | 61,061 | 1,720 | 72,264 |
| | B-cells | 451 | 970 | - | 3,694 | 5,115 |
| | BM | - | - | - | - | **512,208** |
| P3-BM | Mono | 11,425 | 31,655 | - | 24,347 | 67,427 |
| | Neu | 7,687 | 129,203 | 224,483 | 113,752 | 475,125 |
| | Ery | 4,384 | 3,049 | 11,344 | 9,060 | 27,837 |
| | B-cells | 737 | 3,593 | - | 22,894 | 27,224 |
| | BM | - | - | - | - | **597,613** |

Table 1: Cell populations in P1/2/3 bone marrow datasets.

| Cell Population | Cell Counts |
|---|---|
| HSC | 1,334 |
| Early Progenitor (Early Prog) | 3,014 |
| Intermediate Progenitor (Interm Prog) | 257 |
| Late Progenitor (Late Prog) | 256 |
| GMP | 366 |
| MEP | 975 |
| Pro Erythroblast | 223 |
| Basophilic Erythroblast | 610 |
| Polychromatophilic Erythroblast (Polychr. Erythrocytes) | 2,379 |
| Monoblast (Mb) | 1,202 |
| Pro-monocyte | 45 |
| Monocyte | 3,860 |
| Myeloid | 28,198 |
| cDCs | 5,820 |
| Platelet | 3,543 |
| Promyelo/myelocyte (Pro-myelocyte) | 2,288 |
| Neutrophils | 80,709 |
| Pre pDCs | 4,128 |
| pDCs | 2,671 |
| Basophils | 2,025 |
| Double Negative T (Double Neg. T) | 1,718 |
| CD4+ Naïve (CD4+ Naïve T) | 37,930 |
| CD4+ Memory (CD4 Mem T) | 33,170 |
| CD8+ Naïve (CD8+Naïve T) | 35,609 |
| CD8+ Memory (CD8 Mem T) | 32,347 |
| NKT | 3,869 |
| NK | 35,749 |
| Pro B | 343 |
| Pre B I | 439 |
| Pre B II | 725 |
| Early Immature B (Early B) | 3,947 |
| Immature B (Immat B) | 16,977 |
| Plasma Cells | 441 |
| Total | **348,263** |

Table 2: Cell populations in Kimmey 6814 bone marrow dataset.

| Cell Population | Cell Counts |
|---|---|
| HSC | 1,921 |
| Early Progenitor (Early Prog) | 4,152 |
| Intermediate Progenitor (Interm Prog) | 406 |
| Late Progenitor (Late Prog) | 413 |
| GMP | 781 |
| MEP | 951 |
| Pro Erythroblast | 600 |
| Basophilic Erythroblast | 1,155 |
| Polychromatophilic Erythroblast (Polychr. Erythrocytes) | 1,830 |
| Monoblast (Mb) | 4,285 |
| Pro-monocyte | 672 |
| Monocyte | 18,873 |
| Myeloid | 68,831 |
| cDCs | 6,079 |
| Platelet | 5,919 |
| Promyelo/myelocyte (Pro-myelocyte) | 6,448 |
| Neutrophils | 152,761 |
| Pre pDCs | 5,304 |
| pDCs | 4,707 |
| Basophils | 3,360 |
| Double Negative T (Double Neg. T) | 6,481 |
| CD4+ Naïve (CD4+ Naïve T) | 38,811 |
| CD4+ Memory (CD4 Mem T) | 54,667 |
| CD8+ Naïve (CD8+Naïve T) | 55,446 |
| CD8+ Memory (CD8 Mem T) | 65,202 |
| NKT | 20,754 |
| NK | 51,667 |
| Pro B | 297 |
| Pre B I | 919 |
| Pre B II | 4,334 |
| Early Immature B (Early B) | 13,071 |
| Immature B (Immat B) | 43,751 |
| Plasma Cells | 2,882 |
| Total | **647,730** |

Table 3: Cell populations in Kimmey 6796 bone marrow dataset.

| Cell Population | Cell Counts |
|---|---|
| unassigned | 85,297 |
| Mature CD4+ T-cells | 13,964 |
| Erythroblasts | 12,030 |
| Naive CD8+ T-cells | 9,564 |
| Mature CD8+ T-cells | 7,821 |
| Mature CD38lo B-cells | 7,796 |
| Naive CD4+ T-cells | 6,987 |
| CD11bhi Monocytes | 6,779 |
| NK cells | 3,864 |
| Megakaryocytes | 3,684 |
| Myelocyte | 3,025 |
| CD11bmid Monocytes | 1,278 |
| Pre-B II cells | 994 |
| CD11b- Monocytes | 912 |
| Mature CD38mid B-cells | 608 |
| Immature B-cells | 502 |
| GMP | 73 |
| Plasmacytoid_DC_cells | 293 |
| HSC | 261 |
| Platelet | 5 |
| Pre-B I cells | 240 |
| CMP | 253 |
| MEP | 194 |
| MPP | 152 |
| Plasma cells | 468 |
| Total | **167,044** |

Table 4: Cell populations in Levine 13 bone marrow dataset.

| Cell Population | Cell Counts |
|---|---|
| unassigned | 161,443 |
| CD4 T-cells | 26,366 |
| Monocytes | 21,099 |
| CD8 T-cells | 20,108 |
| Mature B-cells | 16,520 |
| Pre-B-cells | 6,135 |
| CD16 negative NK cells | 3,905 |
| CD34+CD38-CD123- HSPCs | 3,295 |
| CD16 positive NK cells | 2,248 |
| pDCs | 1,238 |
| Basophils | 1,207 |
| CD34+CD38lo HSCs | 916 |
| Pro-B-cells | 513 |
| Plasma B cells | 330 |
| CD34+CD38+CD123+ HSPCs | 304 |
| Total | **265,627** |

Table 5: Cell populations in Levine 32 bone marrow dataset.

| Dataset | Number of cells | Number of markers | Memory TimeFlow 2 (GMM) (MB) | Memory TimeFlow 2 (FlowSOM) (MB) | Runtime TimeFlow 2 (GMM) (min) | Runtime TimeFlow 2 (FlowSOM) (min) |
|---|---|---|---|---|---|---|
| Levine 13 | 167,044 | 13 | 1354 | 903 | 24 | 22 |
| Levine 32 | 265,627 | 32 | - | 497 | 101 | 9 |
| Kimmey 6814 | 348,263 | 32 | 139 | 1667 | 140 | 15 |
| P1 | 499,544 | 20 | 164 | 1676 | 75 | 24 |
| P2 | 512,208 | 20 | 154 | 1616 | 73 | 19 |
| P3 | 597,613 | 20 | 189 | 419 | 85 | 55 |
| Kimmey 6796 | 647,730 | 32 | 368 | 2819 | 130 | 42 |

Table 6: Performance comparison of TimeFlow 2 (GMM) vs TimeFlow 2 (FlowSOM) for lineage detection (clustering and optimal transport plans) across datasets. Methods were tested for 10 segments, 35% segment overlap, and maximum 25 GMM clusters or a 5x5 SOM grid.

### Supplementary Figures

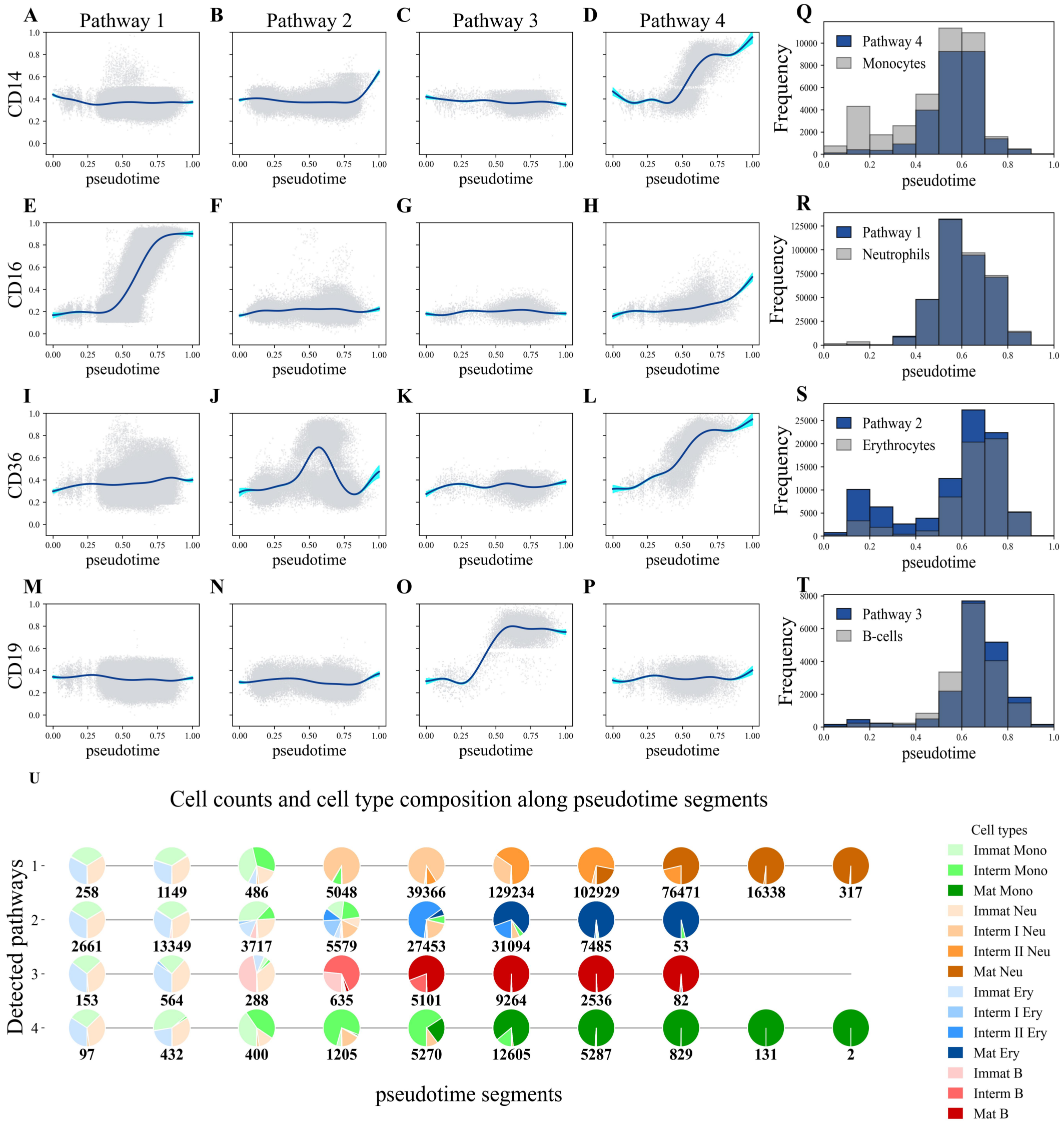


**Supplementary Figure S1**: TimeFlow 2 (FlowSOM) results on the P1-BM dataset. Marker dynamics across the four different pathways/lineages inferred by TimeFlow 2. Each scatterplot dot represents a cell. Both pseudotime and marker expression are scaled in [0,1]. Solid blue curves show the Generalized Additive Model (GAM) fit used to model marker dynamics along pseudotime. Uncertainty around the estimated curve of each pathway is shown with shaded 95% confidence bands. (A-D) GAM fit of CD14 expression across the four pathways. (E-H) GAM fit of CD16 expression across the four pathways. (I-L) GAM fit of CD36 expression across the four pathways. (M-P) GAM fit of CD19 expression across the four pathways. (Q-T) Histograms with overlaid cell distributions along pseudotime. Dark blue histograms represent cell distributions in the inferred pathways, while grey histograms reflect cell distribution of the known lineages based on the P1-BM ground truth gating labels. U) Pie charts showing the counts and cell type composition for each inferred pathway across ten pseudotime segments. Pie chart slices are colored based on the ground truth gating labels, with the total amount of cell counts noted for each pie chart.

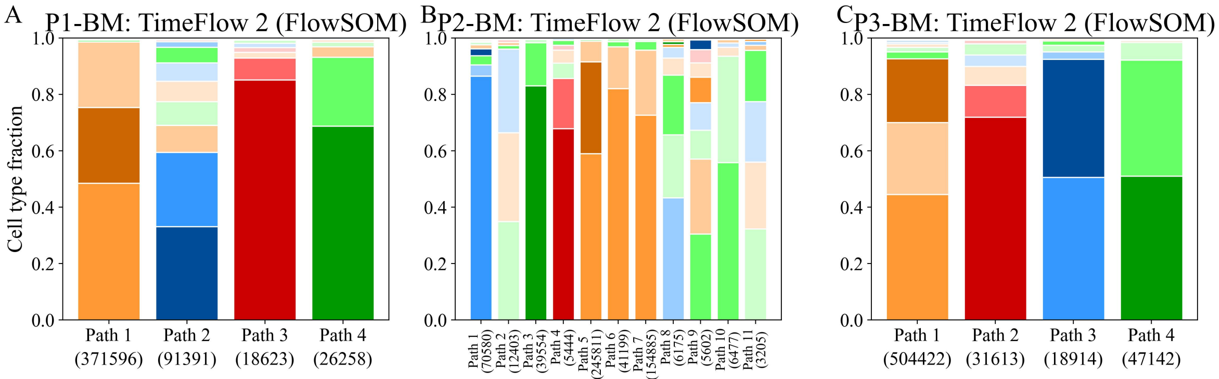


**Supplementary Figure S2**: Side-by-side comparisons of cell type fractions across the different lineages identified by TimeFlow 2 (FlowSOM) in the P1/2/3-BM datasets. (A-C) Stacked bar plots show the cell distribution (%) in each inferred lineage path. The total number of cells assigned to each path is also given. Within each bar, cell types are presented by decreasing proportion. Cell types with larger percentages appear at the base of the bar, while those with smaller percentages appear at the top.

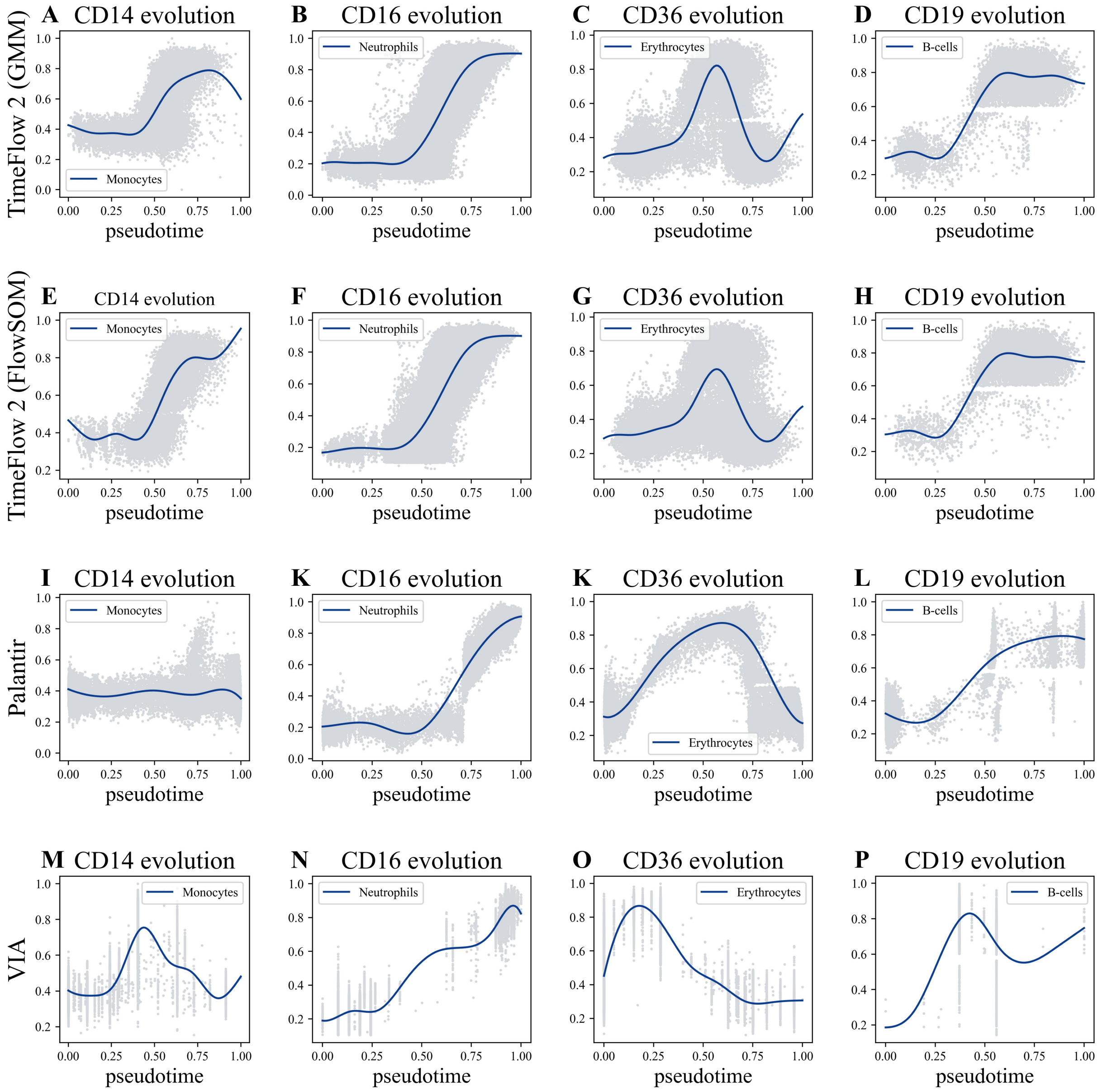


**Supplementary Figure S3**: Comparisons of marker dynamics across the different lineages identified by TimeFlow 2 (GMM), TimeFlow 2 (FlowSOM), Palantir, and VIA in the P1-BM dataset. None of the methods used gating labels during lineage inference. Labels were only used afterwards to retrieve lineages with most mature monocytes, neutrophils, and erythrocytes. Both pseudotime and marker expression are scaled in [0,1]. Each dot in the represents a cell. The solid blue curve shows the Generalized Additive Model (GAM) fit used to model marker dynamics along pseudotime. CD14, CD16, CD36, and CD19 are used for the inferred lineages of monocytes, neutrophils, erythrocytes, and B-cells, respectively. (A-D) Results for TimeFlow 2 (GMM). (E-H) Results for TimeFlow 2 (FlowSOM). (I-L) Results for Palantir. (M-P) Results for VIA.

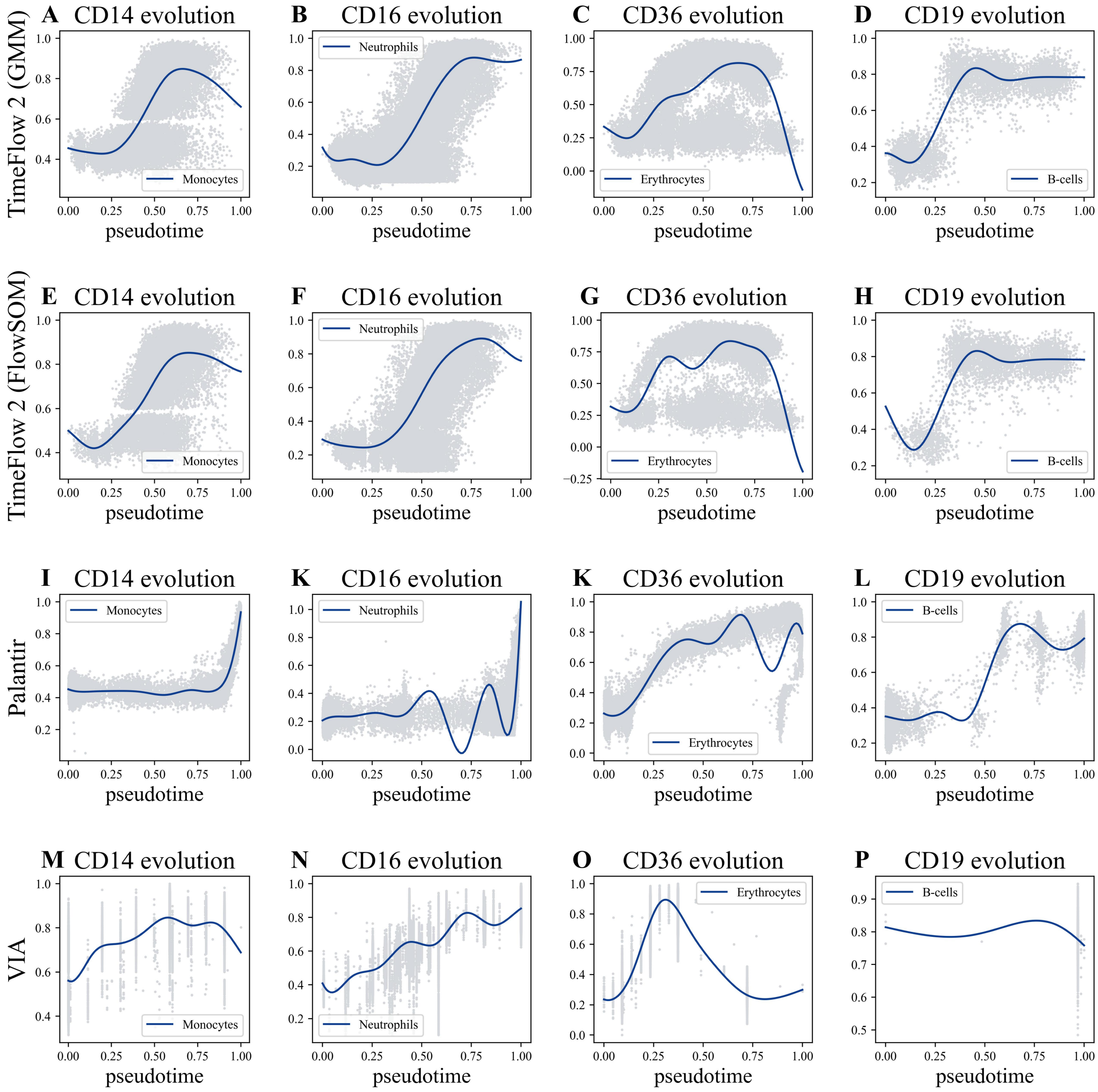


**Supplementary Figure S4**: Comparisons of marker dynamics across the different lineages identified by TimeFlow 2 (GMM), TimeFlow 2 (FlowSOM), Palantir, and VIA in the P2-BM dataset. None of the methods used gating labels during lineage inference. Labels were only used afterwards to retrieve lineages with most mature monocytes, neutrophils, and erythrocytes. Both pseudotime and marker expression are scaled in [0,1]. Each dot in the represents a cell. The solid blue curve shows the Generalized Additive Model (GAM) fit used to model marker dynamics along pseudotime. CD14, CD16, CD36, and CD19 are used for the inferred lineages of monocytes, neutrophils, erythrocytes, and B-cells, respectively. (A-D) Results for TimeFlow 2 (GMM). (E-H) Results for TimeFlow 2 (FlowSOM). (I-L) Results for Palantir. (M-P) Results for VIA.

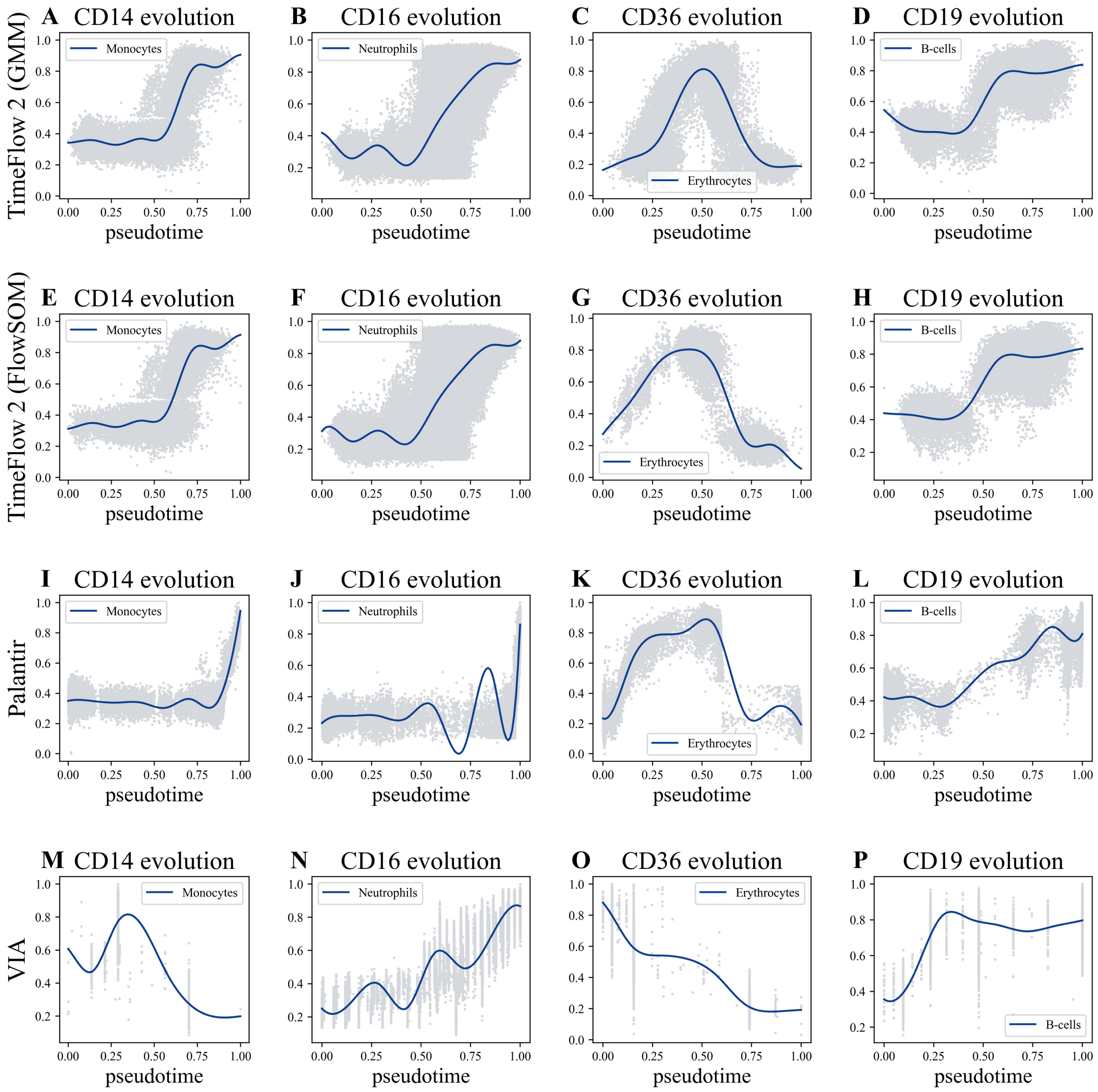


**Supplementary Figure S5**: Comparisons of marker dynamics across the different lineages identified by TimeFlow 2 (GMM), TimeFlow 2 (FlowSOM), Palantir, and VIA in the P3-BM dataset. None of the methods used gating labels during lineage inference. Labels were only used afterwards to retrieve lineages with most mature monocytes, neutrophils, and erythrocytes. Both pseudotime and marker expression are scaled in [0,1]. Each dot in the represents a cell. The solid blue curve shows the Generalized Additive Model (GAM) fit used to model marker dynamics along pseudotime. CD14, CD16, CD36, and CD19 are used for the inferred lineages of monocytes, neutrophils, erythrocytes, and B-cells, respectively. (A-D) Results for TimeFlow 2 (GMM). (E-H) Results for TimeFlow 2 (FlowSOM). (I-L) Results for Palantir. (M-P) Results for VIA.

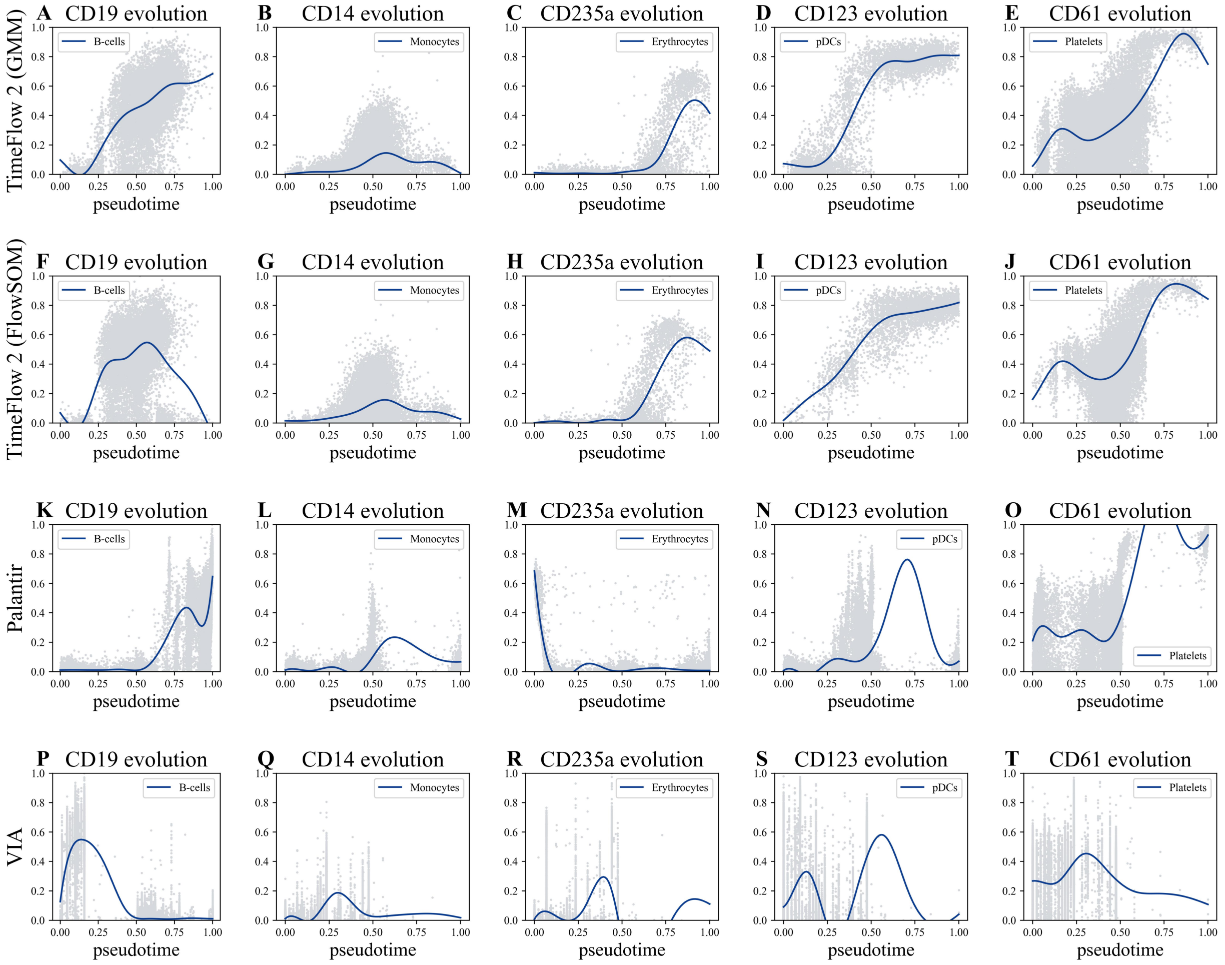


**Supplementary Figure S6**: Comparisons of marker dynamics across the different lineages identified by TimeFlow 2, Palantir, and VIA in the Kimmey 6814 bone marrow dataset. None of the methods used gating labels during lineage inference. Labels were only used afterwards to retrieve lineages with most mature B-cells and monocytes. Both pseudotime and marker expression are scaled in [0,1]. Each dot in the represents a cell. The solid blue curve shows the Generalized Additive Model (GAM) fit used to model marker dynamics along pseudotime. CD19, CD14, CD235a, CD123, and CD61 are used for the inferred lineages of B-cells, monocytes, erythrocytes, pDCs, and platelets, respectively. (A-E) Results for TimeFlow 2 (GMM). (F-J) Results for TimeFlow 2 (FlowSOM). (K-O) Results for Palantir (P-T) Results for VIA.

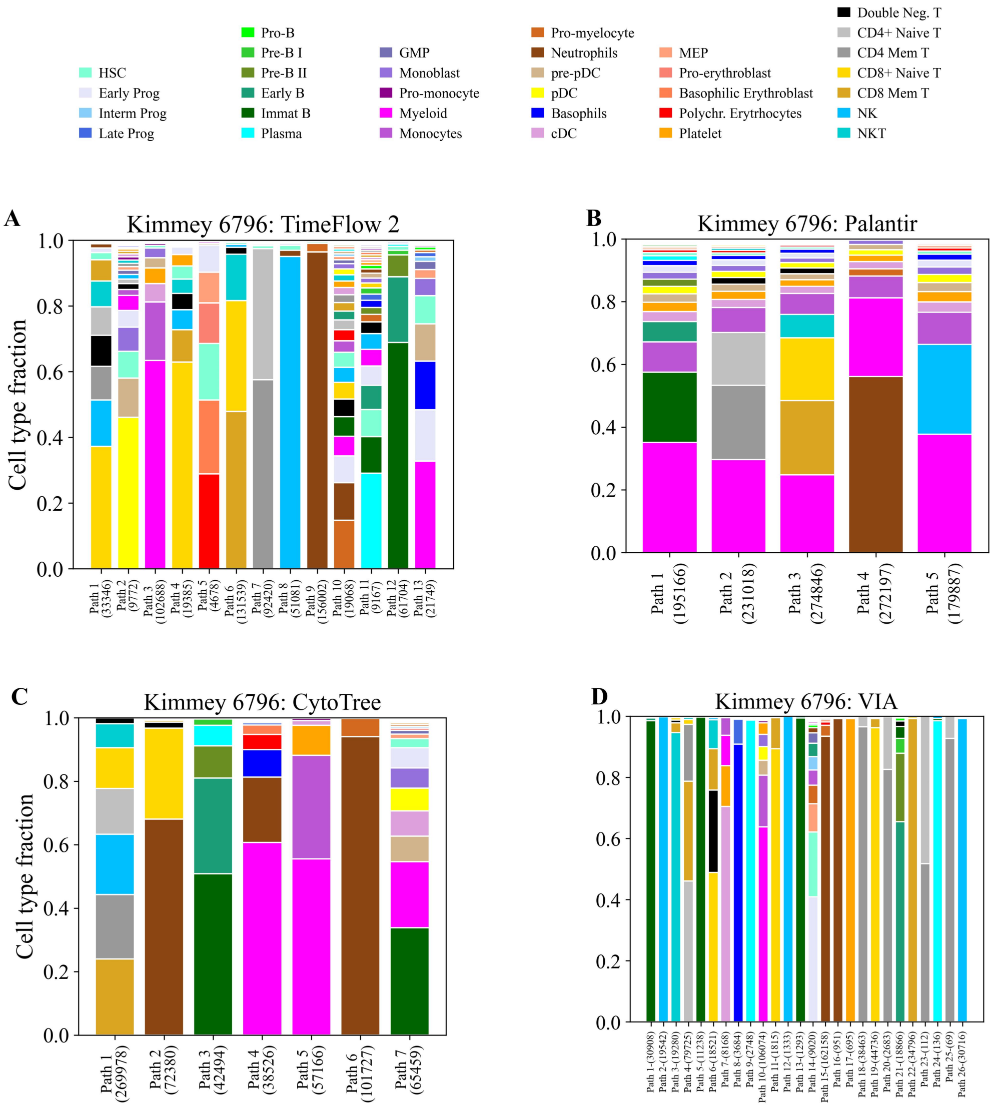


**Supplementary Figure S7**: Side-by-side comparisons of cell type fractions across the different lineages identified by TimeFlow 2, Palantir, CytoTree, and VIA in the Kimmey 6796 bone marrow dataset. Stacked bar plots show the cell distribution (%) in each inferred lineage path. The total number of cells assigned to each path is also given. Within each bar, cell types are presented by decreasing proportion. Cell types with larger percentages appear at the base of the bar, while those with smaller percentages appear at the top. (A) Bar plot for TimeFlow. (B) Bar plot for Palantir. (C) Bar plot for CytoTree. (D) Bar plot for VIA.

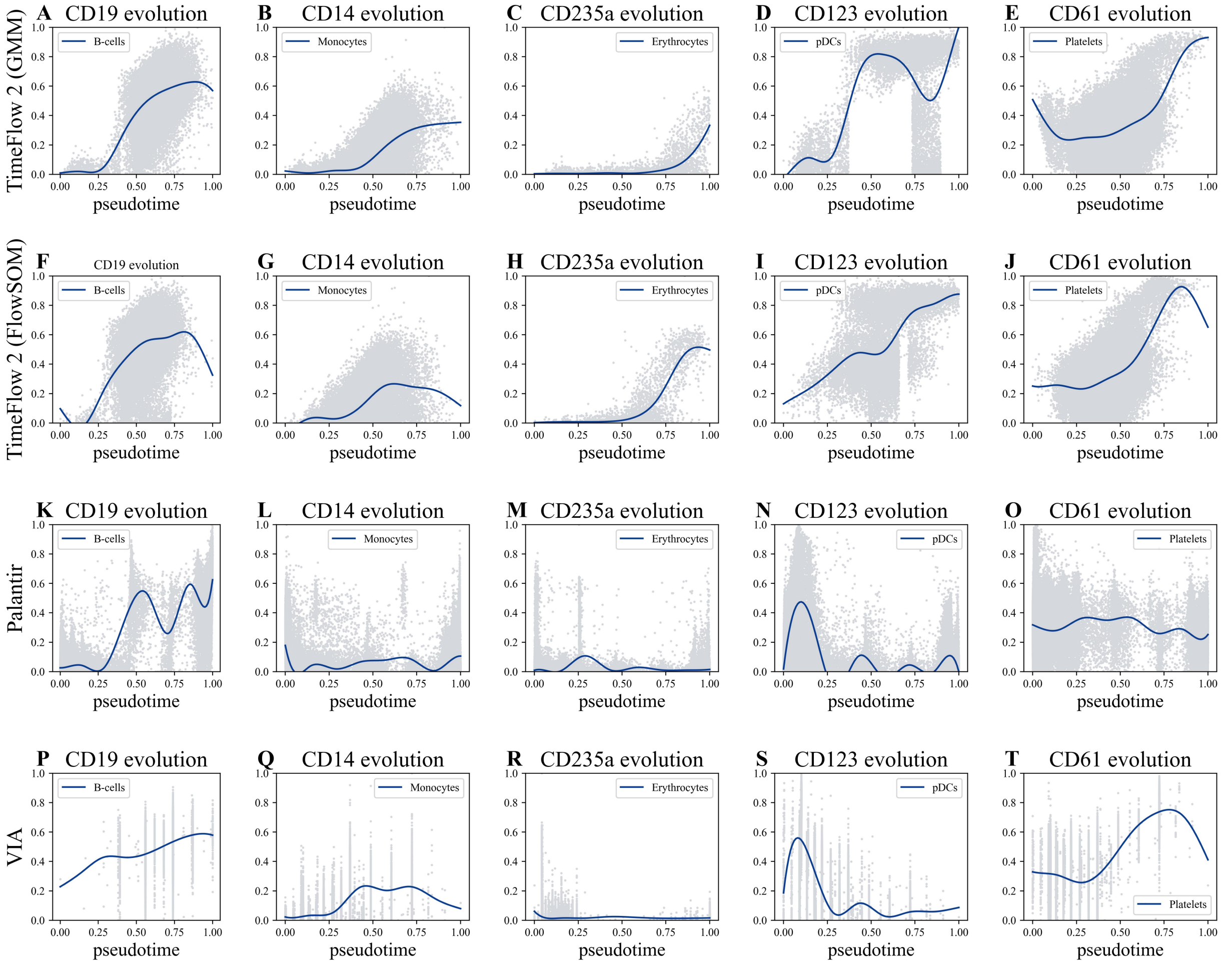


**Supplementary Figure S8**: Comparisons of marker dynamics across the different lineages identified by TimeFlow 2, Palantir, and VIA in the Kimmey 6796 bone marrow dataset. None of the methods used gating labels during lineage inference. Labels were only used afterwards to retrieve lineages with most mature B-cells and monocytes. Both pseudotime and marker expression are scaled in [0,1]. Each dot in the represents a cell. The solid blue curve shows the Generalized Additive Model (GAM) fit used to model marker dynamics along pseudotime. CD19, CD14, CD235a, CD123, and CD61 are used for the inferred lineages of B-cells, monocytes, erythrocytes, pDCs, and platelets, respectively. (A-E) Results for TimeFlow 2 (GMM). (F-J) Results for TimeFlow 2 (FlowSOM). (K-O) Results for Palantir. (P-T) Results for VIA.

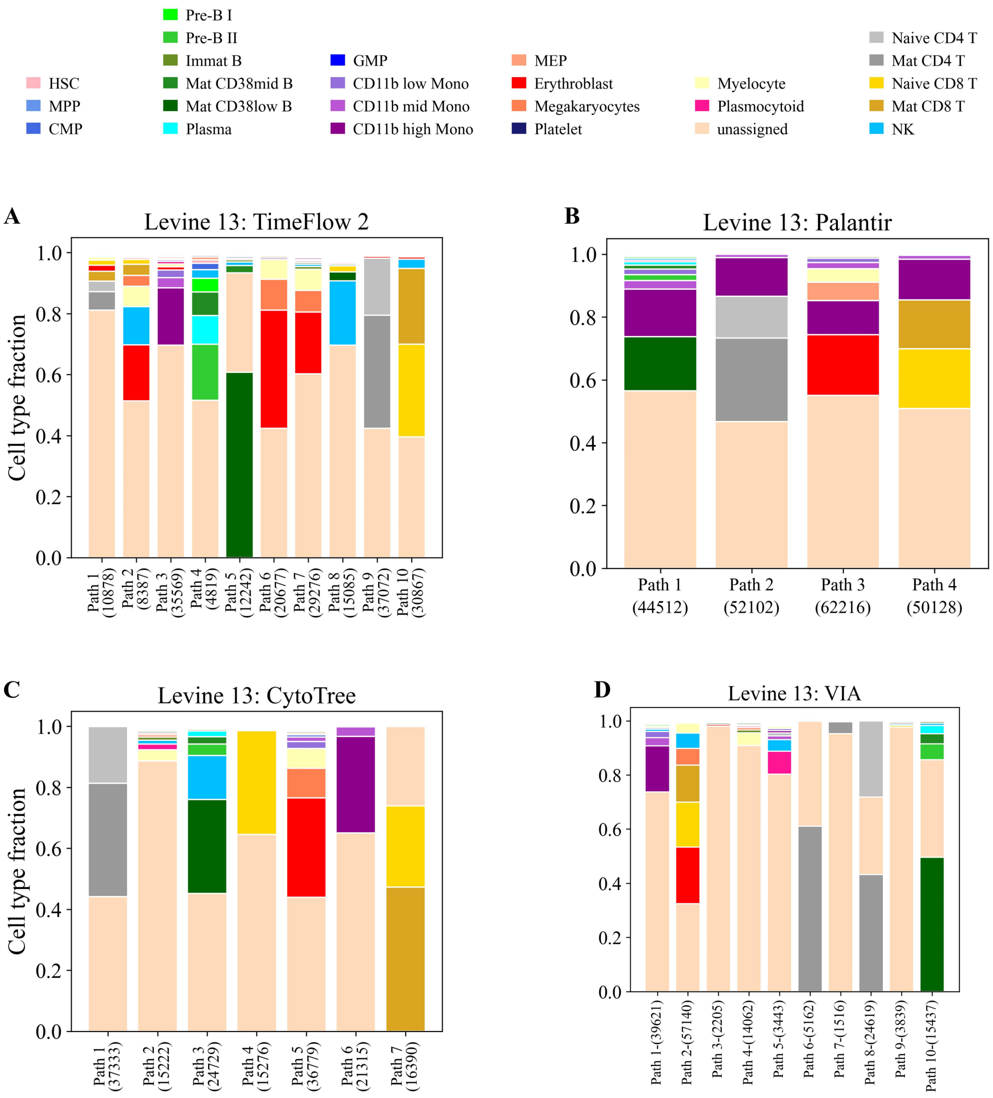


**Supplementary Figure S9**: Side-by-side comparisons of cell type fractions across the different lineages identified by TimeFlow 2, Palantir, CytoTree, and VIA in the Levine 13 bone marrow dataset. Stacked bar plots show the cell distribution (%) in each inferred lineage path. The total number of cells assigned to each path is also given. Within each bar, cell types are presented by decreasing proportion. Cell types with larger percentages appear at the base of the bar, while those with smaller percentages appear at the top. (A) Bar plot for TimeFlow. (B) Bar plot for Palantir. (C) Bar plot for CytoTree. (D) Bar plot for VIA.

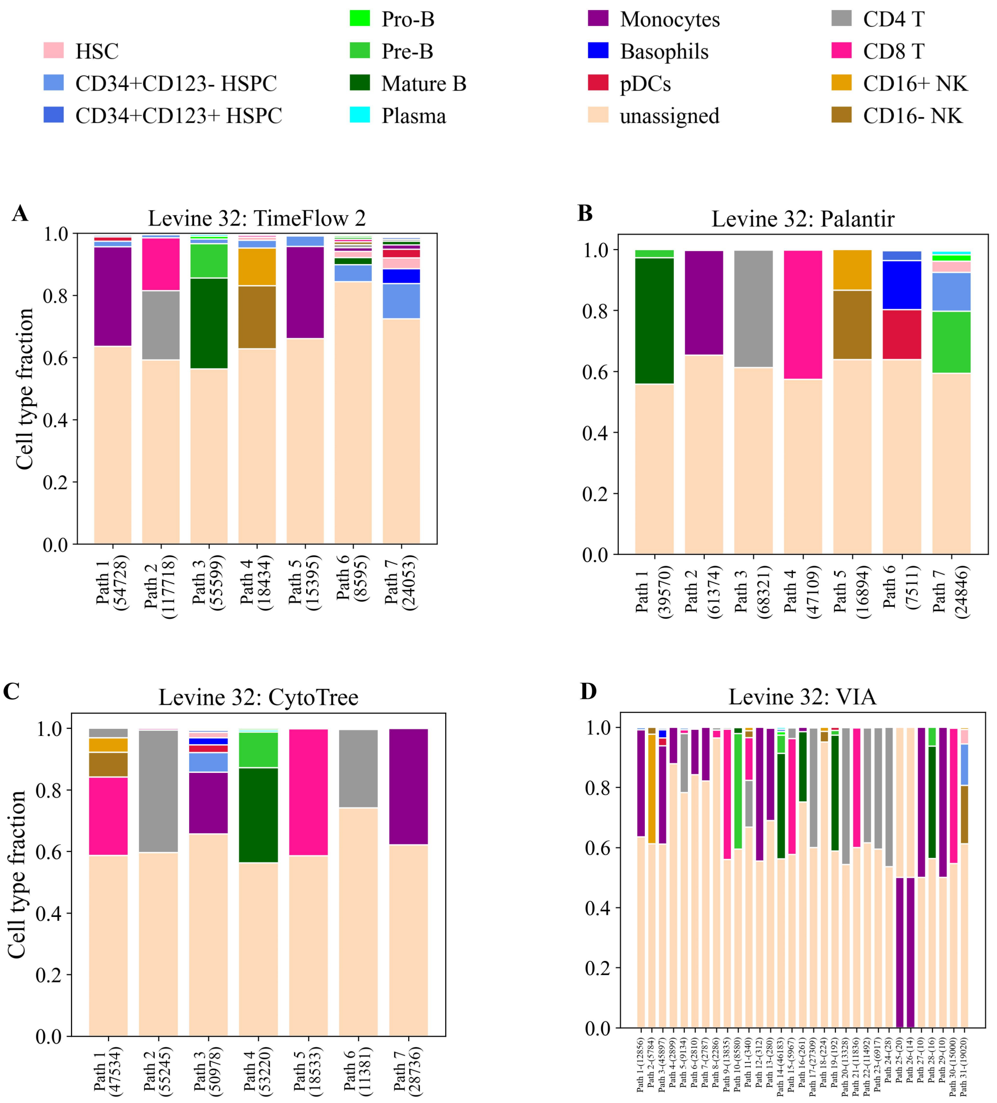


**Supplementary Figure S10**: Side-by-side comparisons of cell type fractions across the different lineages identified by TimeFlow 2, Palantir, CytoTree, and VIA in the Levine 32 bone marrow dataset. Stacked bar plots show the cell distribution (%) in each inferred lineage path. The total number of cells assigned to each path is also given. Within each bar, cell types are presented by decreasing proportion. Cell types with larger percentages appear at the base of the bar, while those with smaller percentages appear at the top. (A) Bar plot for TimeFlow. (B) Bar plot for Palantir. (C) Bar plot for CytoTree. (D) Bar plot for VIA.

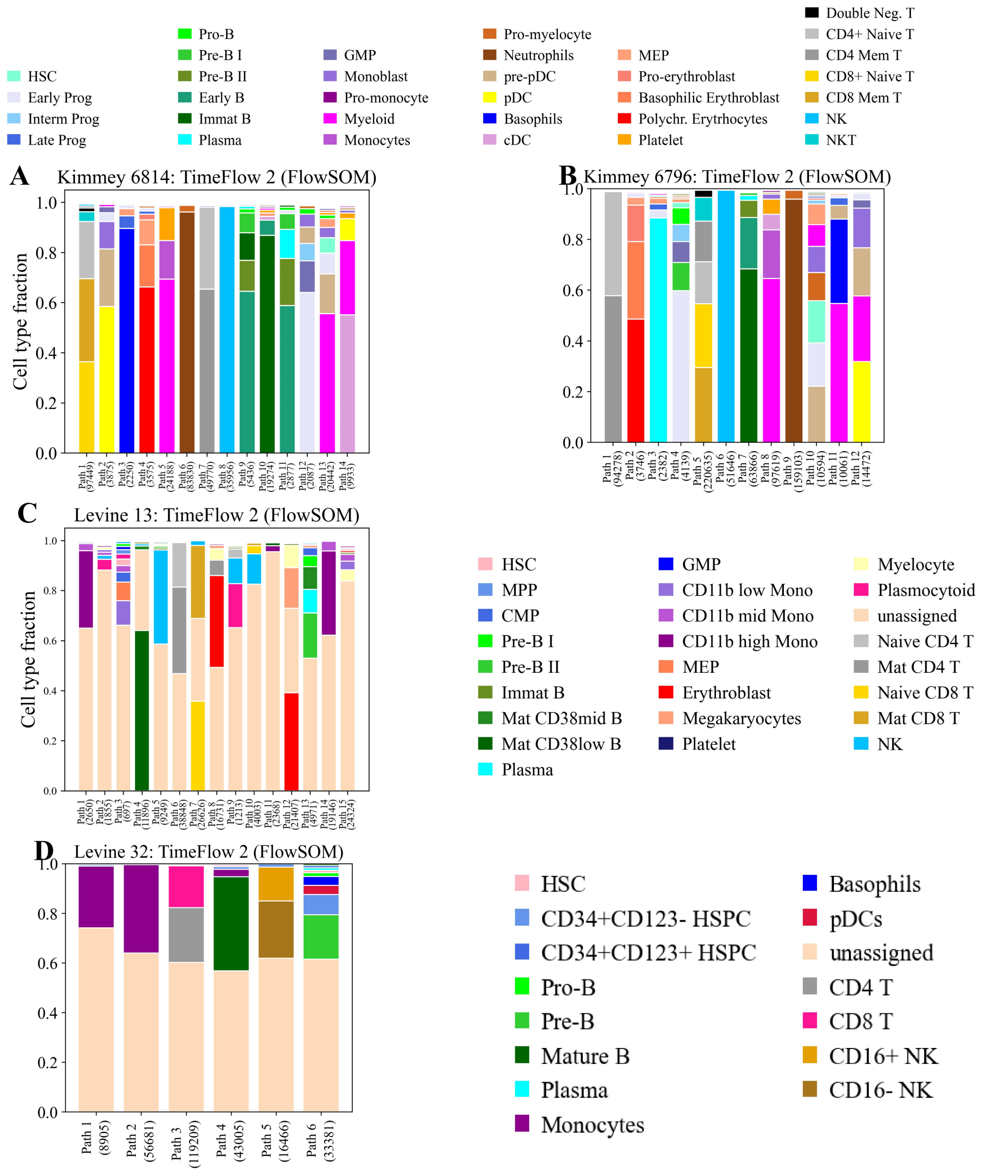


**Supplementary Figure S11**: Comparisons of cell type fractions across the different lineages identified by TimeFlow 2 (FlowSOM) for bone marrow datasets. Stacked bar plots show the cell distribution (%) in each inferred lineage path. The total number of cells assigned to each path is also given. Within each bar, cell types are presented by decreasing proportion. Cell types with larger percentages appear at the base of the bar, while those with smaller percentages appear at the top. (A) Bar plots for Kimmey 6814. (B) Bar plots for Kimmey 6796. (C) Bar plots for Levine 13. (D) Bar plots for Levine 32.

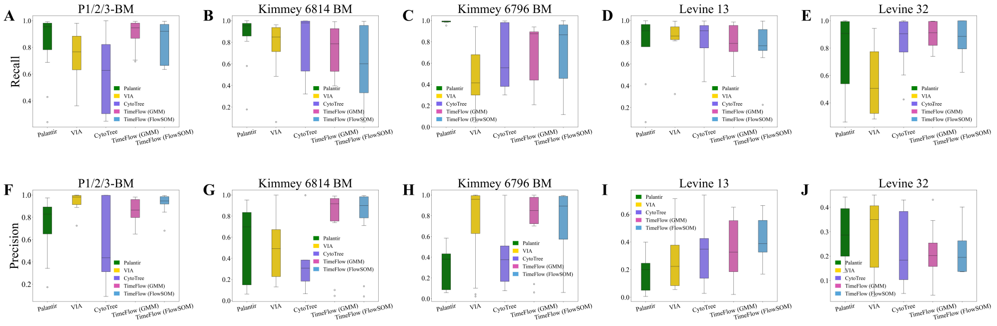


**Supplementary Figure S12**: Methods comparisons across different datasets based on their recall and precision scores for all ground truth lineages identified in a dataset. Boxplots show the median, quartiles, minimum/maximum values, and outliers represented as individual points. (A-E) Results for the recall metric. (F-J) Results for the precision metric. Ground truth lineages in each dataset are described in Section 2.5 and Supplementary Section S3.

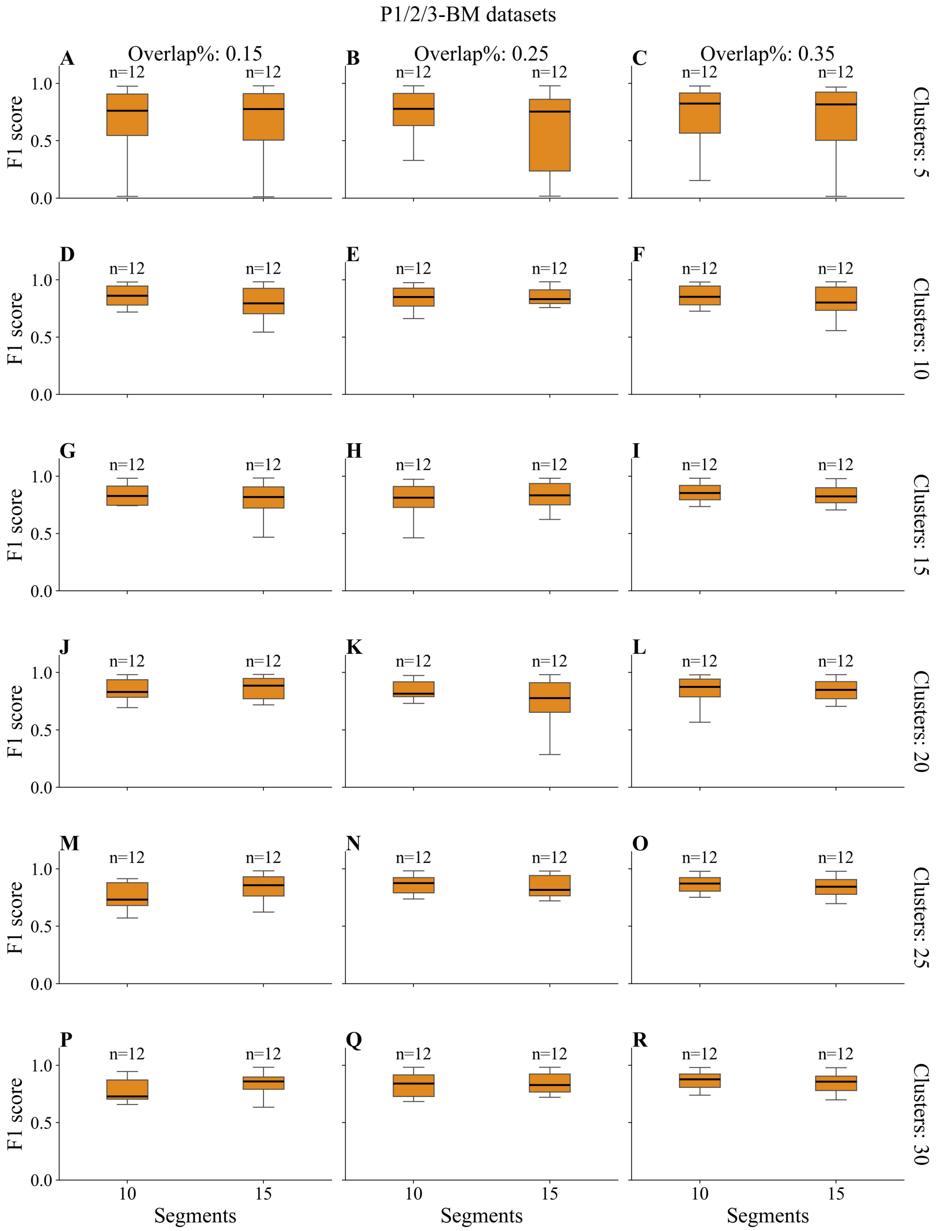


**Supplementary Figure S13**: F1-score sensitivity to hyper-parameter settings on the P1/2/3-BM datasets using TimeFlow 2 (GMM). Each row in the grid corresponds to the number of maximum clusters the Gaussian Mixture Model is allowed to use within each segment, while each column to a different percentage of segment width overlap. For each unique combination of maximum clusters and width overlap, F1-scores are given in cases that TimeFlow 2 correctly detected four lineages in any of the P1/2/3-BM datasets, using either 10 or 15 pseudotime segments. Boxplots show the median, quartiles, minimum/maximum values, and outliers represented as individual points. The number of data points used for each boxplot is denoted by n. Cases with n=0 imply that TimeFlow 2 failed to identify four lineages in any dataset for that configuration, while n=4, n=8, n=12, suggest that TimeFlow detected four lineages in one, two and three patients, respectively.

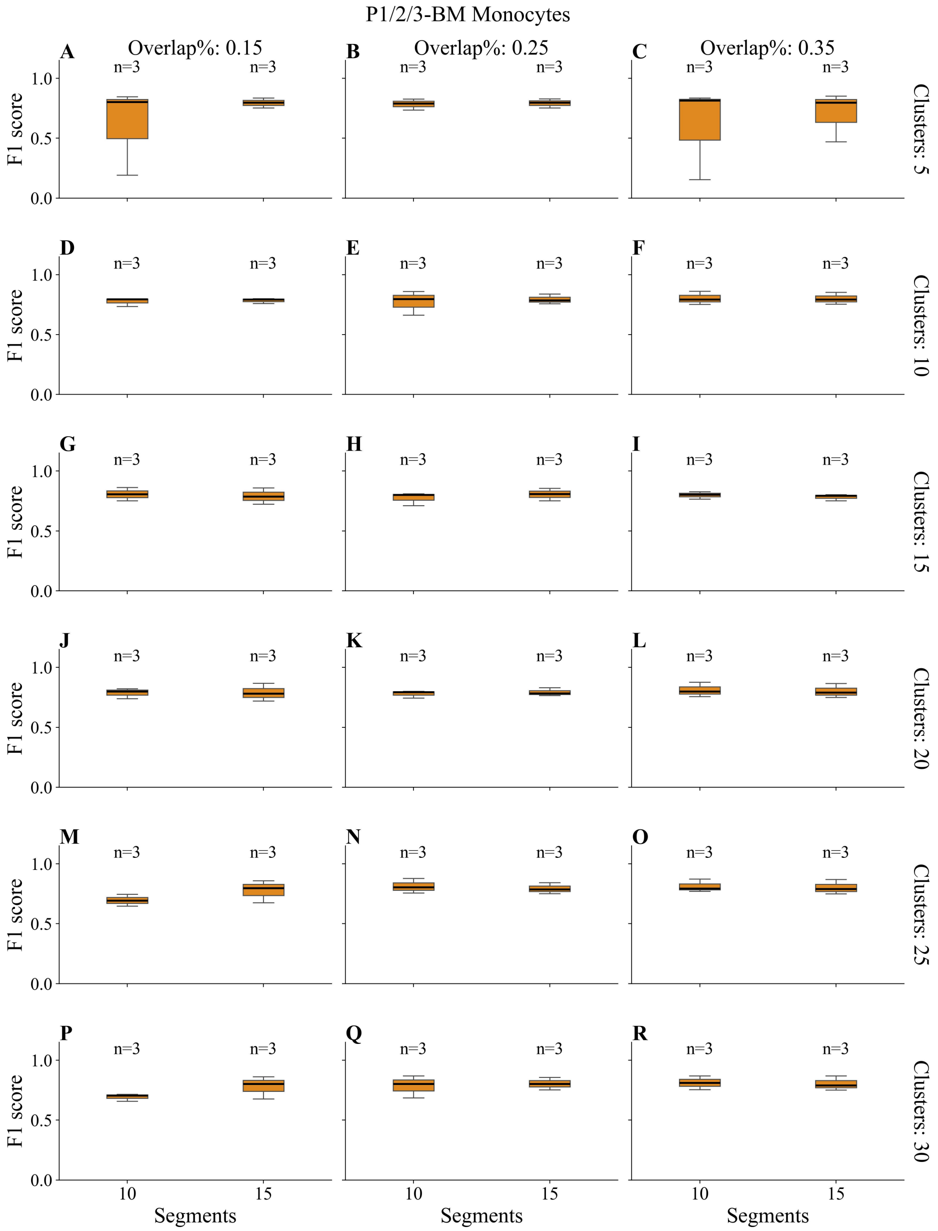


**Supplementary Figure S14**: F1-score sensitivity to hyper-parameter settings only for lineages of monocytes in P1/2/3-BM using TimeFlow 2 (GMM). Each row in the grid corresponds to the number of maximum clusters the Gaussian Mixture Model is allowed to use within each segment, while each column to a different percentage of segment width overlap. For each unique combination of maximum clusters and width overlap, F1-scores are given only for the lineages of monocytes, under the condition that TimeFlow 2 correctly detected four lineages in any of the P1/2/3-BM datasets, using either 10 or 15 pseudotime segments. Boxplots show the median, quartiles, minimum/maximum values, and outliers represented as individual points. The number of data points used for each boxplot is denoted by n. Cases with n=0 imply that TimeFlow 2 failed to identify four lineages in any dataset for that configuration, while n=1, n=2, n=3, suggest that TimeFlow detected four lineages in one, two and three patients, respectively.

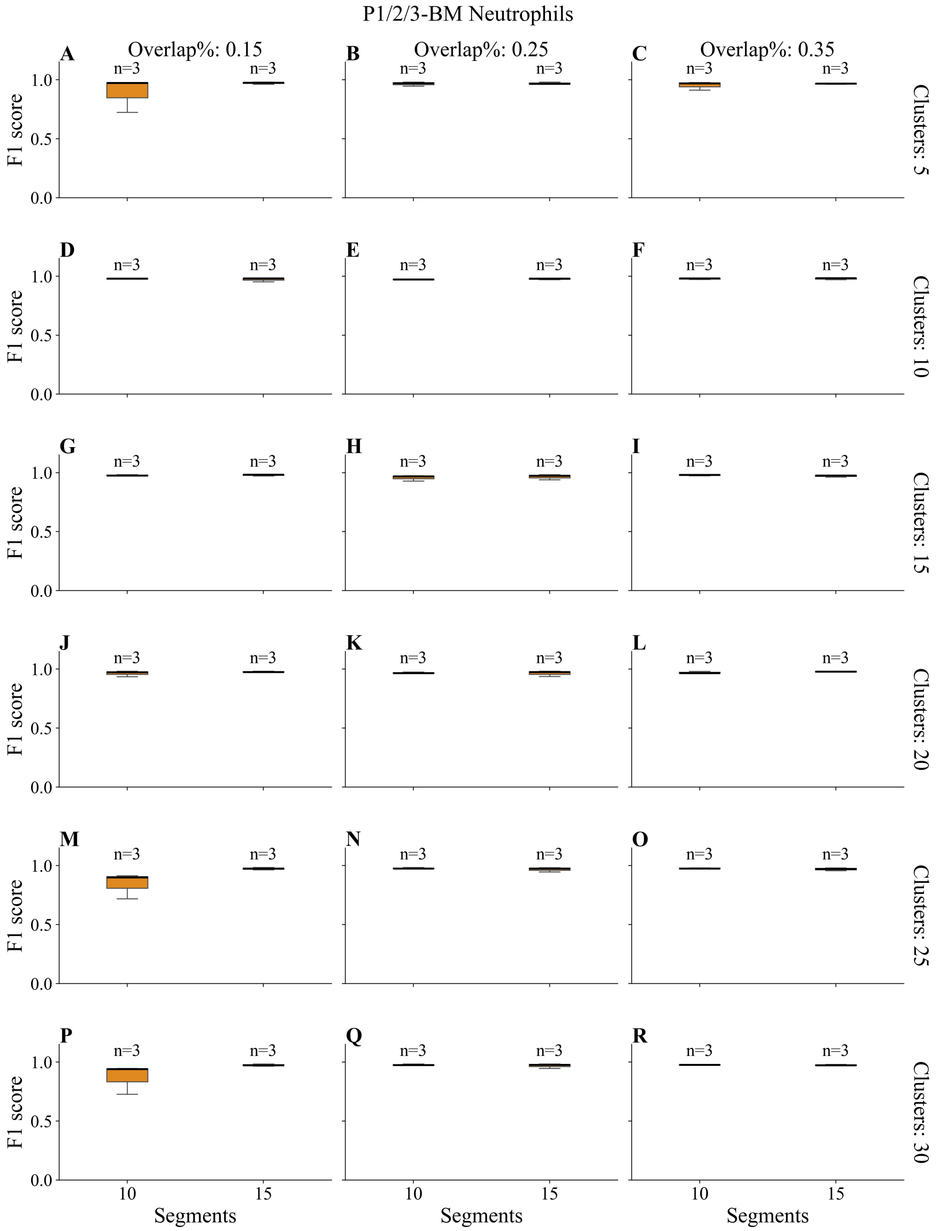


**Supplementary Figure S15**: F1-score sensitivity to hyper-parameter settings only for lineages of neutrophils in P1/2/3-BM using TimeFlow 2 (GMM). Each row in the grid corresponds to the number of maximum clusters the Gaussian Mixture Model is allowed to use within each segment, while each column to a different percentage of segment width overlap. For each unique combination of maximum clusters and width overlap, F1-scores are given only for the lineages of neutrophils, under the condition that TimeFlow 2 correctly detected four lineages in any of the P1/2/3-BM datasets, using either 10 or 15 pseudotime segments. Boxplots show the median, quartiles, minimum/maximum values, and outliers represented as individual points. The number of data points used for each boxplot is denoted by n. Cases with n=0 imply that TimeFlow 2 failed to identify four lineages in any dataset for that configuration, while n=1, n=2, n=3, suggest that TimeFlow detected four lineages in one, two and three patients, respectively.

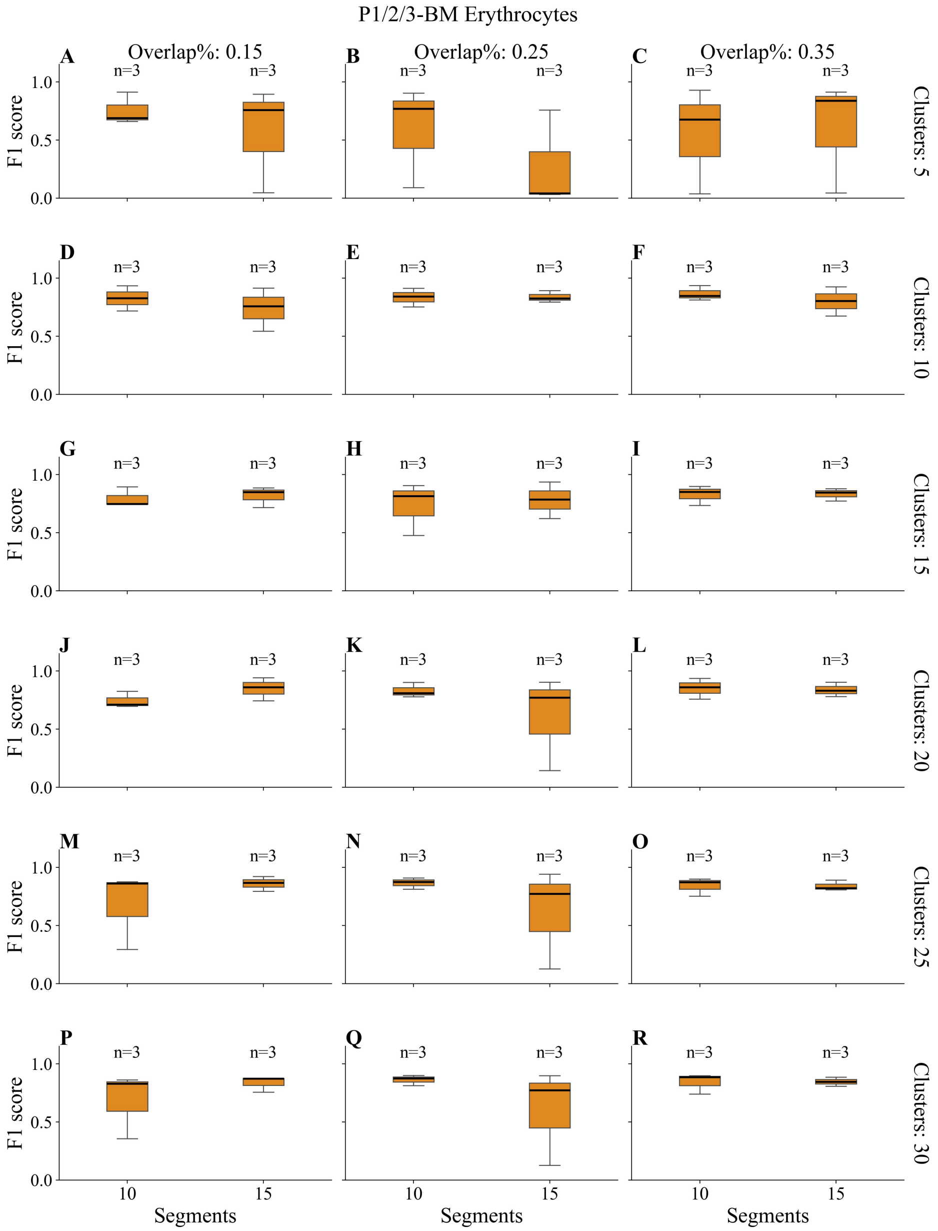


**Supplementary Figure S16**: F1-score sensitivity to hyper-parameter settings only for lineages of erythrocytes in P1/2/3-BM using TimeFlow 2 (GMM). Each row in the grid corresponds to the number of maximum clusters the Gaussian Mixture Model is allowed to use within each segment, while each column to a different percentage of segment width overlap. For each unique combination of maximum clusters and width overlap, F1-scores are given only for the lineages of erythrocytes, under the condition that TimeFlow 2 correctly detected four lineages in any of the P1/2/3-BM datasets, using either 10 or 15 pseudotime segments. Boxplots show the median, quartiles, minimum/maximum values, and outliers represented as individual points. The number of data points used for each boxplot is denoted by n. Cases with n=0 imply that TimeFlow 2 failed to identify four lineages in any dataset for that configuration, while n=1, n=2, n=3, suggest that TimeFlow detected four lineages in one, two and three patients, respectively.

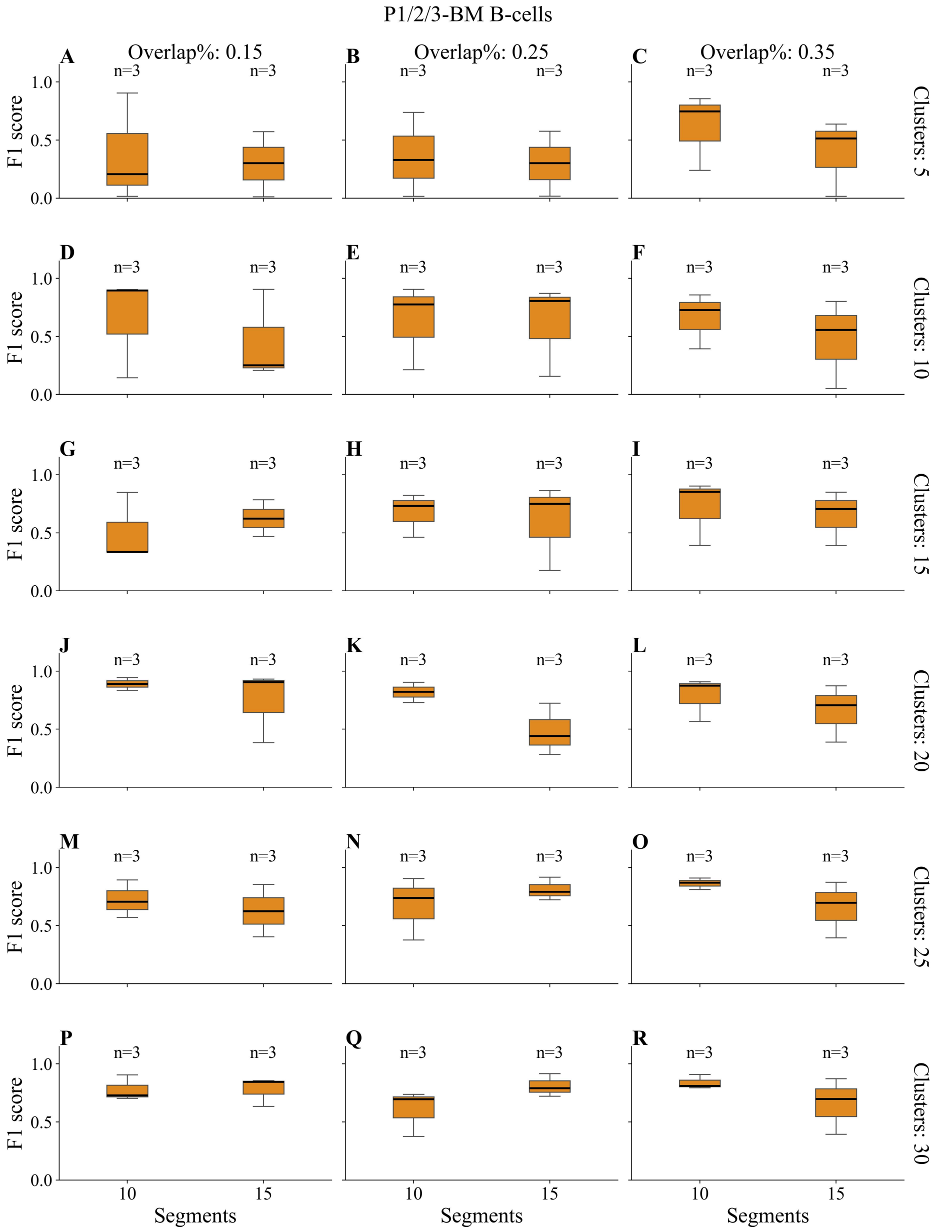


**Supplementary Figure S17**: F1-score sensitivity to hyper-parameter settings only for lineages of B-cells in P1/2/3-BM using TimeFlow 2 (GMM). Each row in the grid corresponds to the number of maximum clusters the Gaussian Mixture Model is allowed to use within each segment, while each column to a different percentage of segment width overlap. For each unique combination of maximum clusters and width overlap, F1-scores are given only for the lineages of B-cells, under the condition that TimeFlow 2 correctly detected four lineages in any of the P1/2/3-BM datasets, using either 10 or 15 pseudotime segments. Boxplots show the median, quartiles, minimum/maximum values, and outliers represented as individual points. The number of data points used for each boxplot is denoted by n. Cases with n=0 imply that TimeFlow 2 failed to identify four lineages in any dataset for that configuration, while n=1, n=2, n=3, suggest that TimeFlow detected four lineages in one, two and three patients, respectively.

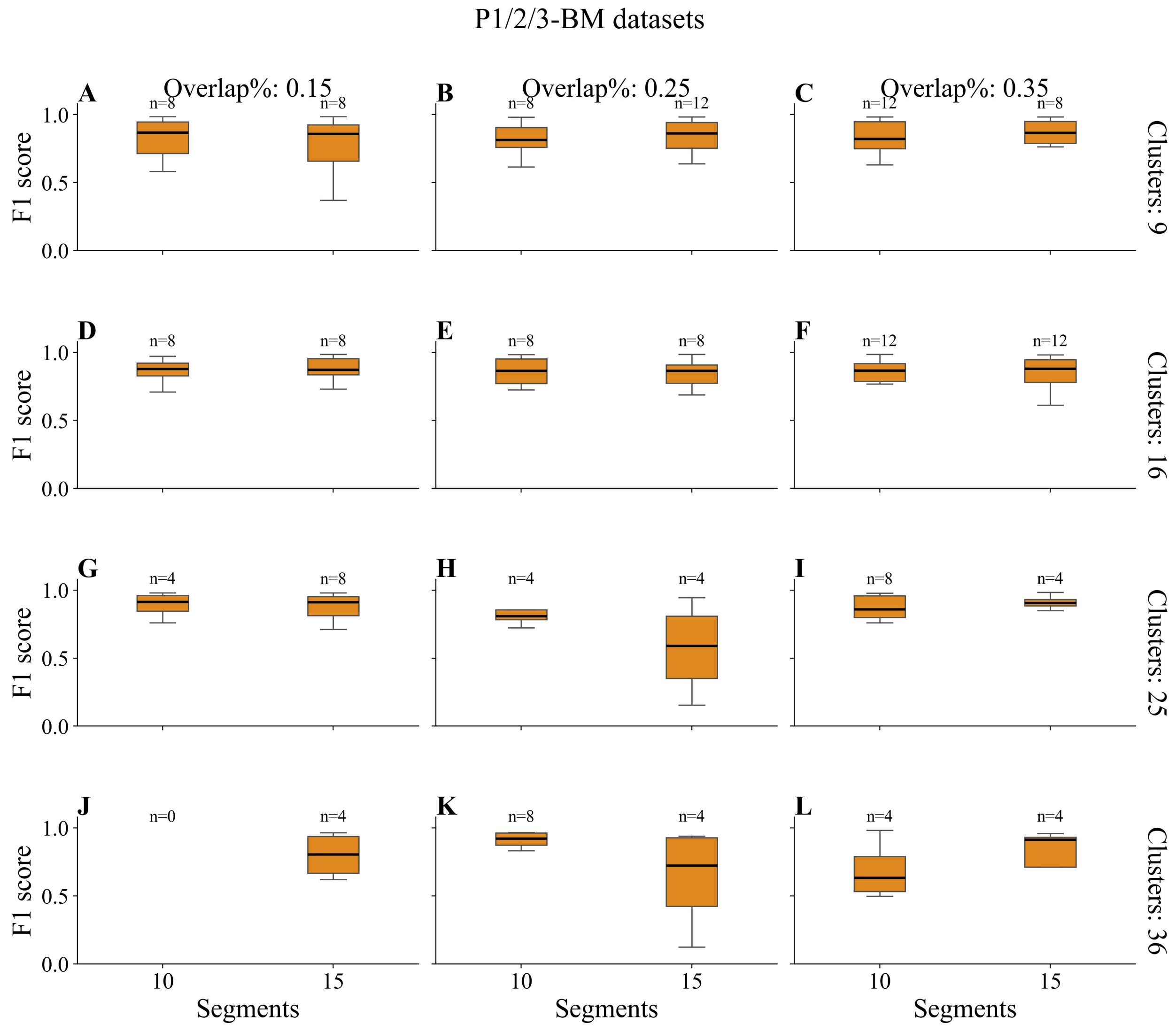


**Supplementary Figure S18**: F1-score sensitivity to hyper-parameter settings on the P1/2/3-BM datasets using TimeFlow 2 (FlowSOM). Each row in the grid corresponds to the number of SOM nodes in each segment, while each column to a different percentage of segment width overlap. For each unique combination of maximum clusters and width overlap, F1-scores are given in cases that TimeFlow 2 correctly detected four lineages in any of the P1/2/3-BM datasets, using either 10 or 15 pseudotime segments. Boxplots show the median, quartiles, minimum/maximum values, and outliers represented as individual points. The number of data points used for each boxplot is denoted by n. Cases with n=0 imply that TimeFlow 2 failed to identify four lineages in any dataset for that configuration, while n=4, n=8, n=12, suggest that TimeFlow detected four lineages in one, two and three patients, respectively.

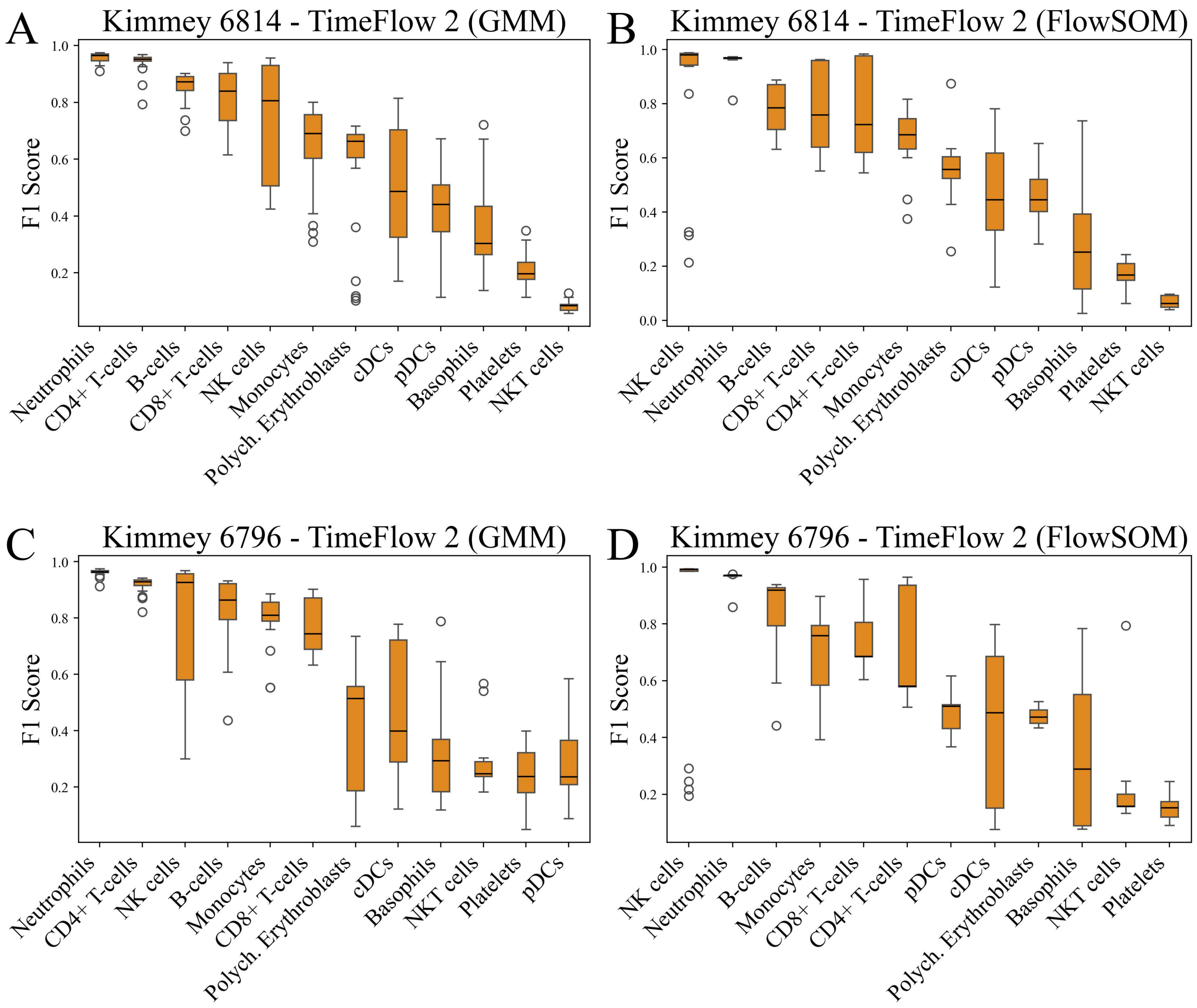


**Supplementary Figure S19**: F1-score sensitivity to hyper-parameter settings for Kimmey 6814 and Kimmey 6796. Comparisons between TimeFlow 2 (GMM) and TimeFlow 2 (FlowSOM) across different cell populations and datasets based on F1 score. Boxplots show the median, quartiles, minimum/maximum values, and outliers represented as individual points. (A-B) Results for Kimmey 6814. (C-D) Results for Kimmey 6796.

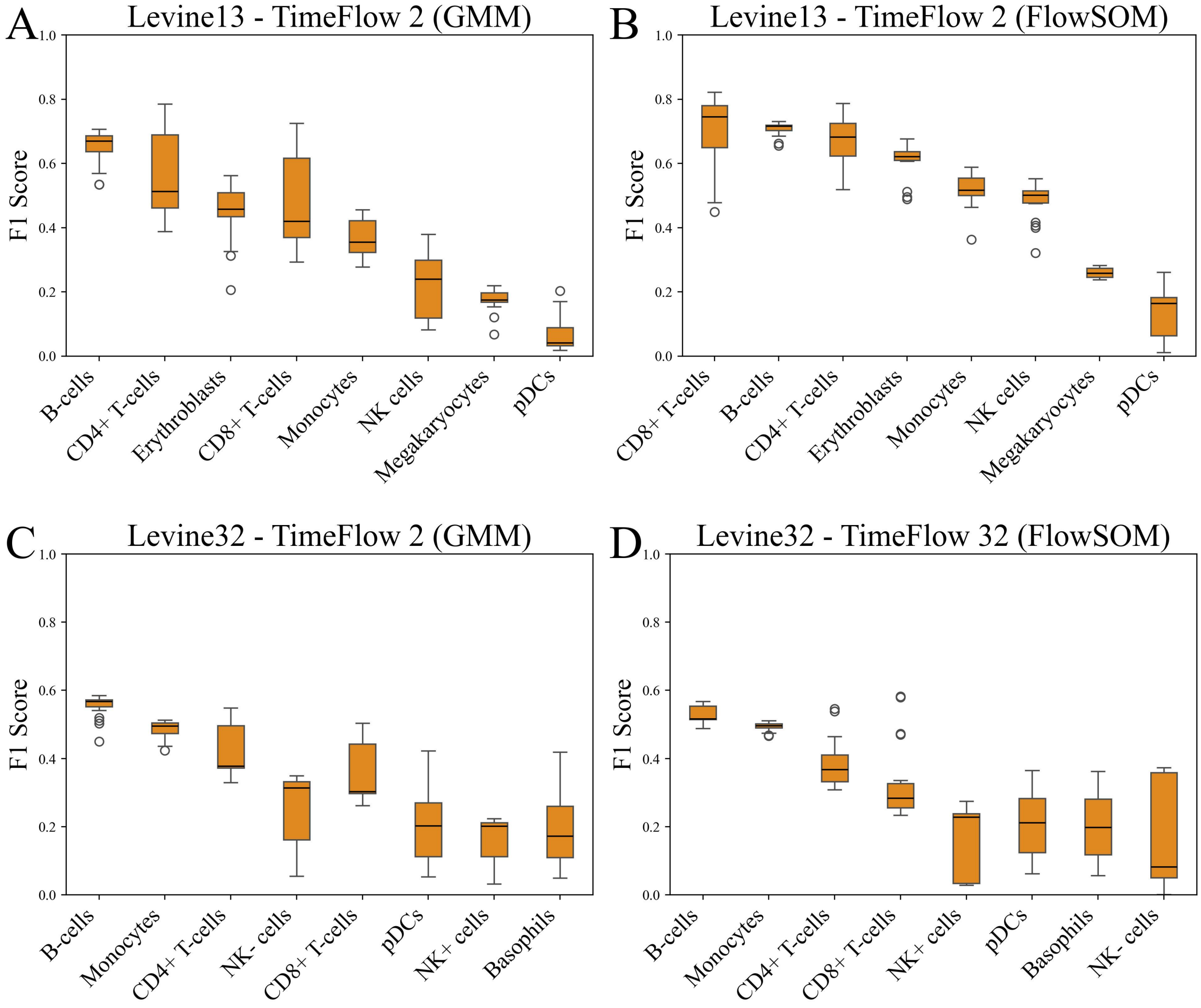


**Supplementary Figure S20**: F1-score sensitivity to hyper-parameter settings for Levine 13 and Levine 32. Comparisons between TimeFlow 2 (GMM) and TimeFlow 2 (FlowSOM) across different cell populations and datasets based on F1 score. Boxplots show the median, quartiles, minimum/maximum values, and outliers represented as individual points. (A-B) Results for Levine 13. (C-D) Results for Levine 32.

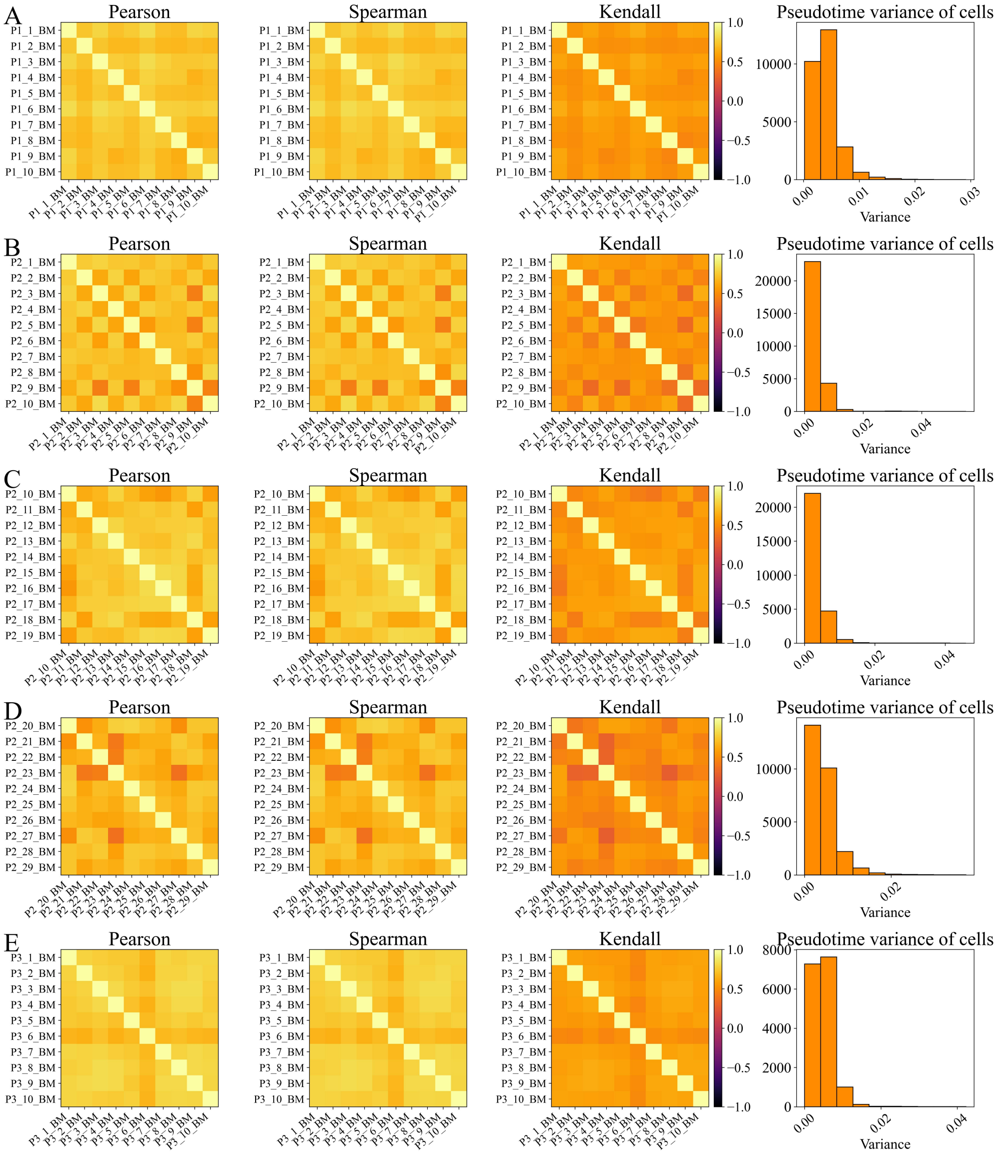


**Supplementary Figure S21**: Assessment of pseudotime uncertainty for random subsets of cells: Pearson correlation heatmap for pseudotime values, Spearman and Kendall correlation heatmaps for cell rankings, and histogram of cells' pseudotime variance. (A) Results for subsets of P1-BM. (B) Results for subsets of P2.1-BM cohort. (C) Results for subsets of P2.2-BM cohort. (D) Results for subsets of P2.3-BM cohort. (E) Results for subsets of P3-BM.

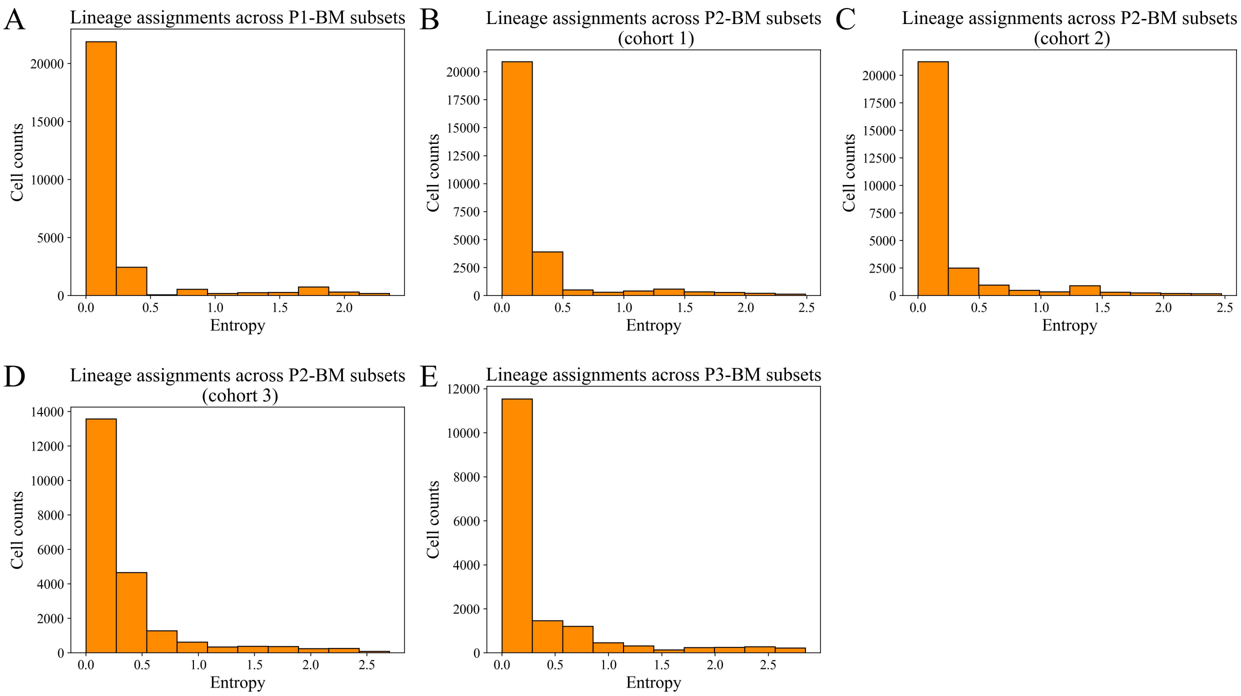


**Supplementary Figure S22**: Assessment of lineage assignment consistency across random subsets based on Shannon entropy. (A) Histogram of entropy scores for subsets of P1-BM. (B) Histogram of entropy scores for subsets of P2.1-BM cohort. (C) Histogram of entropy scores for subsets of P2.2-BM cohort. (D) Histogram of entropy scores for subsets of P2.3-BM cohort. (E) Histogram of entropy scores for subsets of P3-BM.

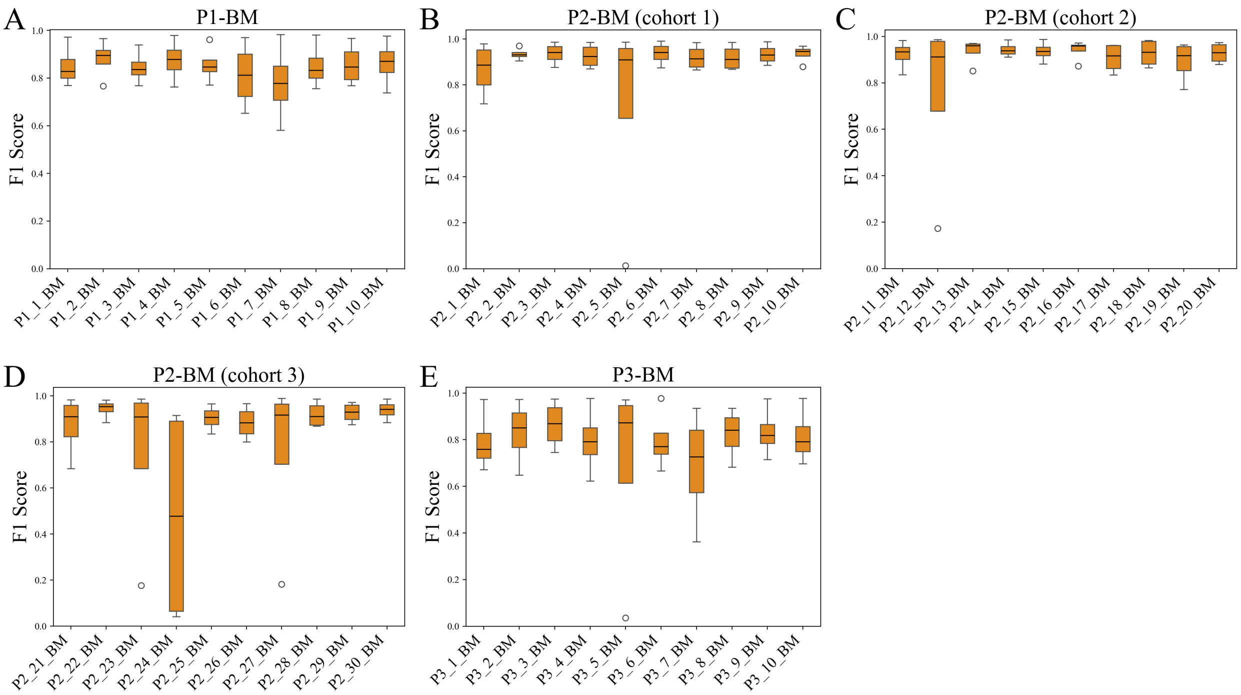


**Supplementary Figure S23**: Assessment of lineage detection across random subsets based on F1 score. (A) Results for subsets of P1-BM. (B) Results for subsets of P2.1-BM cohort. (C) Results for subsets of P2.2-BM cohort. (D) Results for subsets of P2.3-BM cohort. (E) Results for subsets of P3-BM. Boxplots show the median, quartiles, minimum/maximum values, and outliers represented as individual points.

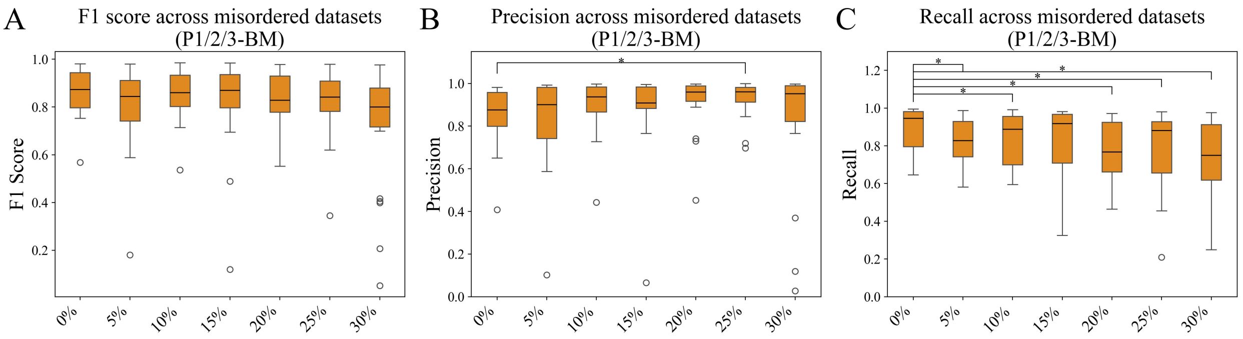


**Supplementary Figure S24**: Assessment of lineage detection across datasets with distorted pseudotime for different proportions based on F1 score. (A) F1 score for P1/2/3-BM datasets with misordered cells. (B) Precision for P1/2/3-BM datasets with misordered cells. (C) Recall for P1/2/3-BM datasets with misordered cells. Boxplots show the median, quartiles, minimum/maximum values, and outliers represented as individual points. Benjamini & Hochberg adjusted p values correspond to paired one-sided t tests with a significance level of 0.05 (*signifies p value $< 0.05$, **: p value $< 0.01$, ***: p value $< 0.001$).
